# Neural Dynamics of the Processing of Speech Features: Evidence for a Progression of Features from Acoustic to Sentential Processing

**DOI:** 10.1101/2024.02.02.578603

**Authors:** I.M Dushyanthi Karunathilake, Christian Brodbeck, Shohini Bhattasali, Philip Resnik, Jonathan Z. Simon

## Abstract

When we listen to speech, our brain’s neurophysiological responses “track” its acoustic features, but it is less well understood how these auditory responses are enhanced by linguistic content. Here, we recorded magnetoencephalography (MEG) responses while subjects listened to four types of continuous-speech-like passages: speech-envelope modulated noise, English-like non-words, scrambled words, and a narrative passage. Temporal response function (TRF) analysis provides strong neural evidence for the emergent features of speech processing in cortex, from acoustics to higher-level linguistics, as incremental steps in neural speech processing. Critically, we show a stepwise hierarchical progression of progressively higher order features over time, reflected in both bottom-up (early) and top-down (late) processing stages. Linguistically driven top-down mechanisms take the form of late N400-like responses, suggesting a central role of predictive coding mechanisms at multiple levels. As expected, the neural processing of lower-level acoustic feature responses is bilateral or right lateralized, with left lateralization emerging only for lexical-semantic features. Finally, our results identify potential neural markers, linguistic level late responses, derived from TRF components modulated by linguistic content, suggesting that these markers are indicative of speech *comprehension* rather than mere speech perception.

**Significance Statement:** We investigate neural processing mechanisms as speech evolves from acoustic signals to meaningful language, using stimuli ranging from without any linguistic information to fully well-formed linguistic content. Computational models based on speech and linguistic hierarchy reveal that cortical responses time-lock to emergent features from acoustics to linguistic processes at the sentence level, with increasing the semantic information in the acoustic input. Temporal response functions (TRFs) uncovered millisecond-level processing dynamics as speech and language stages unfold. Each speech feature undergoes early and late processing stages, with the former driven by bottom-up activation and the latter influenced by top-down mechanisms. These insights enhance our understanding of the hierarchical nature of auditory language processing.

## Introduction

Human language is known for its hierarchical structure, and in the course of speech understanding, the brain first performs computations on the acoustic waveform, which further undergo processing through various intermediate stages, integrating both bottom-up and top-down mechanisms (Davis et al., 2011; Arnal et al., 2016). Prior research has shown that these neural processing stages align with at least some levels in the speech and linguistic hierarchy (Gillis et al., 2021; Brodbeck et al., 2022; Keshishian et al., 2023), including acoustic analysis, phonological analysis, lexical processing, and contextual processing. However, the specific temporal dynamics and how these processes emerge during discourse level speech processing are still not well understood. Identifying the neural bases underlying these stages and their roles in bottom-up and top-down mechanisms deepen our understanding of the neural markers that might be utilized to evaluate cognitive processes beyond basic sensory processing, such as intelligibility and semantic processing.

Previous functional magnetic resonance imaging (fMRI) research has shown numerous brain regions that are sensitive to specific aspects of language understanding (Xu et al., 2005; Lerner et al., 2011; Deniz et al., 2023), but the inherently limited temporal resolution of fMRI poses challenges in investigating fast temporal dynamics of speech comprehension. Imaging modalities with higher temporal resolution, magneto/electroencephalography (M/EEG), often employ stimuli of short length (a few seconds or less), leading to a focus on word processing rather than capturing the broader aspects of spoken language (Alday, 2019). Recently, advances in neural speech-tracking measures such as the temporal response function (TRF) paradigm, have allowed investigators to study time-locked neural responses to many different speech features, and in more ecologically valid settings. These neural speech-tracking measures are well established for acoustic properties of the speech (speech envelope) (Ding and Simon, 2012a; Brodbeck et al., 2020). Additional research has revealed that many linguistic elements of speech, sub-lexical, lexical, and context-based properties, also exhibit neural tracking (Brodbeck et al., 2018; Heilbron et al., 2022) above and beyond auditory neural tracking. How these tracking measures depend on the linguistic content of the speech, however, is still poorly understood, e.g., as a function of semantic information available.

To answer these questions, we employed MEG to record the neural responses of subjects listening to four types of speech materials (Figure 1A). In addition to ordinary narrative speech, we also presented word-scrambled narrative speech (with word-level semantic content but no more), narrated non-words (which sounds like speech but with no semantic information whatsoever), and envelope-modulated noise (entirely unintelligible even at the phoneme level). Thus, each passage type was designed to neurally progress through the brain only up to a specific level in the hierarchy of speech processing: acoustic processing (for speech modulated noise), phoneme and word-boundary identification (narrated non-words), word meaning (scrambled narration), and full construction and processing of structured meaning (narrative), respectively. All four stimulus types exhibited similar accent, speech-like prosody, and rhythm across passages.

**Figure 1.**
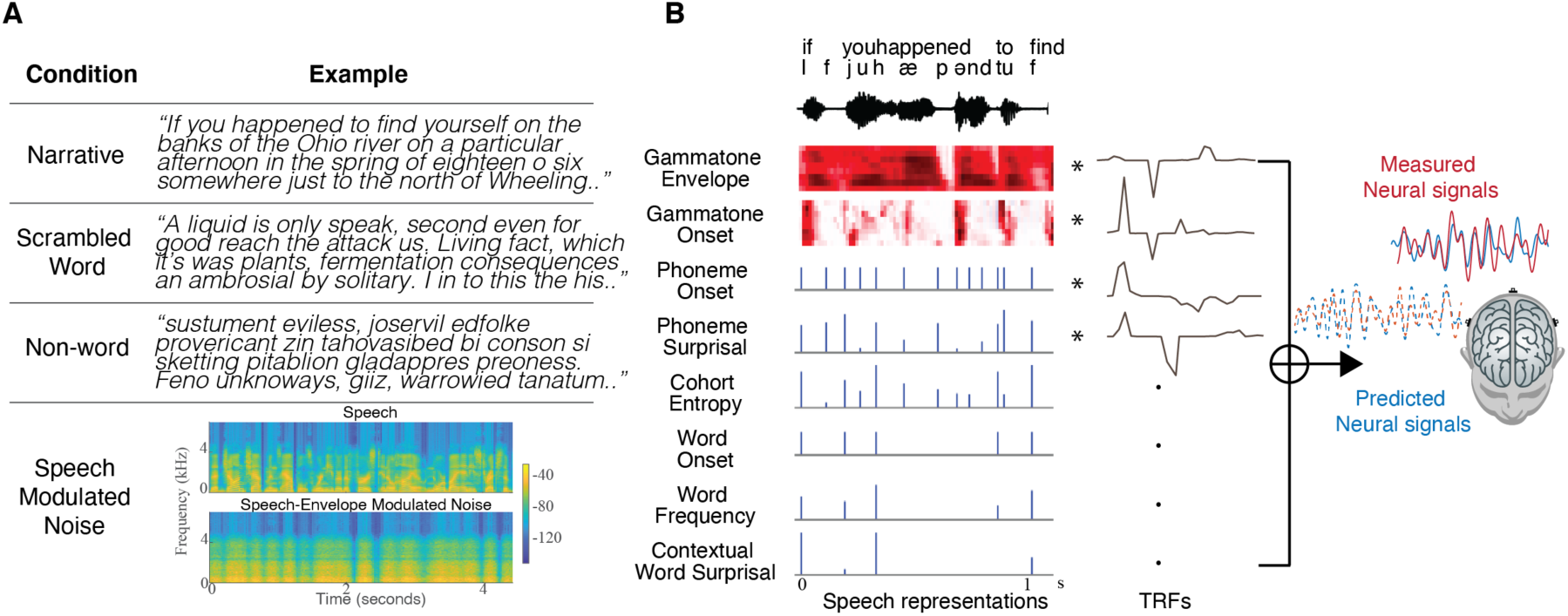
Overview of the study design and analysis framework. (A). Examples of the four stimulus types. Participants listened to 1-minute-long speech passages of each passage type while magnetoencephalography (MEG) brain activity was recorded. All stimuli had similar prosody and rhythm. Speech-modulated noise (bottom) is unintelligible and its spectro-temporal characteristics are shown in the bottom row. (B). Multivariate temporal response functions (mTRFs) were used to model the brain activity at different levels of speech representations and at each current dipole. Orthographic and phonemic transcriptions aligned with a sample acoustic waveform are shown for reference. Speech representations includes acoustic features (8-band auditory gammatone spectrogram; acoustic envelope and acoustic onset), sub-lexical features (phoneme onset, phoneme surprisal and cohort entropy) and lexical features (word onset, word frequency (unigram word surprisal) and contextual word surprisal).

Using multivariate TRF analysis, with different TRFs contributing simultaneously from acoustic to contextual levels, we investigate how different feature representations progress in the brain across different processing levels (Figure 1B). We hypothesize that the ascending brain processing stages will show emergent features, from acoustic to sentence-level linguistic, as incremental steps in the processing of the speech occurs. We additionally expect that many speech features require more than one processing stage: early processing, which is primarily bottom-up, and late processing, which is primarily top-down, consistent with the corrections that may be required for predictive coding models, analogous to generalized N400 event related potential (ERP) responses (Nour Eddine et al., 2024). Further, we anticipate that hemispheric lateralization will vary with passage type and that lower-level processing would generally manifest bilaterally or with a weak right hemisphere advantage, whereas left-lateralization would be predicted for lexico-semantic processing. Lastly, we aim to investigate the progression of the temporal dynamics of speech processing, and its reorganization in response to changes in the linguistic content of the sensory input.

## Materials and Methods

### Participants

34 native English speaking younger adults participated in this experiment. Data from four subjects were excluded from the analysis because of technical issues during data acquisition (1 subject) and poor performance on the behavioral tasks (see experimental procedure) (3 subjects), leaving thirty participants in the analysis (15 females, mean age 22 y, age range 18-29 y, 1 left-handed). All participants reported normal hearing and no history of neurological disorders. All experimental procedures were approved by the Internal Review Board of the University of Maryland, College Park. The participants gave their written informed consent before the experiment and received monetary compensation, or course credit (1 subject).

### Speech stimuli

Four types of speech stimuli: narrative, scrambled word, non-words and speech-modulated noise were generated as described below (sample materials are shown in Figure 1A and can be listened to at https://dushk88.github.io/progression-of-neural-features/). Text used for speech stimuli were excerpts from the book “The Botany of Desire” by Michael Pollan (Pollan, 2001). Speech stimuli were computer synthesized using Google text to speech API (Oord et al., 2016) (gTTS) (see example: https://cloud.google.com/text-to-speech). The use of modern text-to-speech synthesizers provides human-like, natural-sounding speech (Aoki et al., 2022; Herrmann, 2023), and ensures acoustic parameters like speech rate, rhythm, and emphasis are consistent across passage types, which is crucial for comparing neural responses across passage types in the current study (Ding et al., 2017) (Figure 1-1).

The narrative (structured and meaningful) passages were excerpts from the first section of the book. A separate section of the book was used for which the words were randomly permuted to create the scrambled word (intermediate structure) passages. Another section, non-overlapping with the previous passages, was used to generate the speech-modulated noise (unintelligible speech) passages. For the non-word (gibberish) passages, nonsense words were extracted from https://www.soybomb.com/tricks/words/ and were randomly arranged to form a continuous passage. Initial versions of both scrambled and non-word passages lacked punctuation marks, but since silences and pauses between words and sentences create natural sounding and rhythmic speech, and in gTTS pauses and silences are cued by punctuation marks, punctuation marks were manually added to the scrambled and non-word passages (using the distribution of the number of words between punctuation marks in the original book).

Speech was synthesized with gTTS using the English US accent male voice and Google Wavenet voice type “en-US-Wavenet-J” (https://google.com/text-to-speech/docs/voices) at the default sampling rate 24 kHz. Once the speech passages were generated, audio files were lowpass filtered below 4 kHz since the MEG audio delivery (air tube) system has a lowpass cutoff of ∼4 kHz. Then the silence segments were trimmed to 400 ms and the audio stimuli were resampled to 22.5 kHz. For each of the speech stimulus types, 1-minute-duration excerpts were extracted.

For construction of the modulated noise passage, the corresponding speech stimuli generated for modulated noise passages were further modified. First, stationary noise was generated with the same frequency spectrum as the speech by randomizing the phases of the stimulus frequency spectrum and inverting back to the time domain. In order to add back the lost rhythmicity to the noise, the stationary speech shaped noise was then modulated with the corresponding slow speech envelope of the original speech (Figure 1A). The slow speech envelope was extracted by low pass filtering (with a 5 Hz cutoff) the Hilbert envelope of the speech passage.

### Experimental procedure

The experiment was conducted in four blocks. Each block comprised of one passage from each passage type, and each passage was repeated twice. The order of passage types was counterbalanced across subjects. The narrative passages were presented in chronological order to preserve the story line to increase the subjects’ attention. In total, each participant listened to a total of 32 trials (4 blocks × 4 types × 2 repetitions = 32 trials) and 8 trials from each passage type (4 blocks × 2 repetitions), where a trial is defined as a presentation of 1-minute-long stimulus passage. At the start of each passage type, subjects were instructed which passage type they were about to listen to. A probe question (depending on the type of passage) was included for each passage (counting occurrences of a probe word; a contextual question based on the story passage; judging which emotion was conveyed in the speech-modulated noise passage) to help maintain participant’s attention to the listening task. Participants who correctly answered at least 70% of the questions (excluding the emotion judgement) were included in the analysis.

The subjects lay supine during the entire experiment and were asked to minimize body movements. Subjects kept their eyes open and fixated at a center of a grey screen. The stimuli were delivered bilaterally at ∼70 dB SPL with E-A-RTONE 3A tubes (impedance 50 Ω) which severely attenuate frequencies above 3 – 4 kHz, and E-A-RLINK (Etymotic Research, Elk Grove Village, United States) disposable earbuds inserted into ear canals.

### Data acquisition and preprocessing

Neuromagnetic data were recorded inside a dimly lit, magnetically shielded room (Vacuumschmelze GmbH & Co. KG, Hanau, Germany) with a whole head 157-channel MEG system (KIT, Kanazawa, Japan), installed at the Maryland Neuroimaging Center. The data were recorded with a sampling rate of 1 kHz along with an online low-pass filter (< 200 Hz) and a 60 Hz notch filter. Three additional sensor channels were employed as environment reference channels.

All data analyses were performed in mne-python 0.23.0 (Gramfort, 2013; Gramfort et al., 2014) and eelbrain 0.36 (Brodbeck et al., 2023). Flat channels were excluded and the environmental magnetic interference was suppressed using temporal signal space separation (tSSS) (Taulu and Simola, 2006). MEG data were then filtered between 1 and 60 Hz using a zero-phase FIR filter (mne-python 0.23.0 default settings). Artifacts such as ocular, cardiac, and muscle artifacts were reduced using independent component analysis (ICA) (Bell and Sejnowski, 1995). The cleaned data were then low pass filtered at 10 Hz and downsampled to 100 Hz for further analysis.

### Neural source localization

The scalp surface (> 2000 points), five head position indicator (HPI) coils (three placed on the forehead, left and right ear), and anatomical landmarks (nasion, left and right periauricular) of each participant were digitized using Polhemus 3SPACE FASTRAK three-dimensional digitizer. The position of the participant’s head relative to the sensors was determined before and after the experiment using HPI coils attached to the scalp surface and the two measurements were averaged. The digitized head shape and the HPI coils locations were used to co-register the template FreeSurfer “fsaverage” (Fischl, 2012) brain to each participant’s head shape using rotation, translation, and uniform scaling.

A neural source space was generated by four-fold icosahedral subdivision of the white matter surface of the fsaverage brain, with the constraint that all source dipoles be oriented perpendicular to the cortical surface. The source space data and the noise covariance estimated from empty room data were used to compute inverse operator via minimum norm current estimation (Dale and Sereno, 1993; Hämäläinen and Ilmoniemi, 1994). The subsequent analyses were limited to frontal, temporal, and parietal brain regions based on the ‘aparc’ FreeSurfer parcellation (Desikan et al., 2006).

### Predictor variables

The speech signal was analyzed in distinct feature spaces that represent various levels of the language hierarchy. These features were grouped into four primary categories: acoustic properties (i.e., acoustic envelope and acoustic onsets), sub-lexical properties (i.e., phoneme onset, phoneme surprisal, and cohort entropy), lexical properties (i.e., word onset and word frequency), and contextual features (i.e., contextual word surprisal). The methodology for generating each of these predictors is detailed below. Overall, these predictors were generated using a combination of signal processing techniques, automatic speech recognition (ASR) systems, and probabilistic models. All predictor variables were downsampled to 100 Hz.

#### Acoustic features

The acoustic envelope predictor is a measure of the amplitude modulation of the speech signal, and reflects the acoustic power/energy of the speech signal over time. In contrast, the acoustic envelope onset predictor is a measure of the salient transients of the speech signal, which are particularly prominent at the beginning of syllables or phonemes. The acoustic envelope and acoustic onsets were computed based on the human auditory system inspired gammatone filters computed by the Gammatone Filterbank Toolkit 1.0 (Heeris, 2018), using 256 center frequencies with cut-off frequencies ranging logarithmically from 20 to 5000 Hz. Each frequency band’s envelope was resampled to 1000 Hz and transformed to log scale. The resulting envelope spectrogram was then averaged into 8 logarithmically spaced frequency bands to obtain the final acoustic envelope predictor. Eight bands were chosen as a trade-off between computational efficiency and the ability to capture detailed information about the amplitude modulation. The acoustic onset representations were computed using the above gammatone acoustic envelope 256-band spectrogram, by applying an auditory edge detection algorithm (Fishbach et al., 2001; Brodbeck et al., 2023). The onset spectrogram was averaged into the same 8 logarithmically spaced frequency bands as the envelope predictor. The distributions of the acoustic envelope and onset predictors were found to be comparable across speech conditions, non-words, scrambled and narrative passages. However, some variations were observed between the speech stimuli and the speech modulated noise stimuli, as evidenced by the comparisons shown in Figure 1-2A. This discrepancy may be attributed to the diminishment of formants and/or sharp onsets in the non-speech (due to its modulation being induced only by the broad band envelope of the speech stimuli).

#### Phoneme onsets and word onsets

Preliminary speech audio alignment for the occurrence of discrete words and phonemes was accomplished using the Montreal Forced Aligner (McAuliffe et al., 2017). Grapheme to phoneme conversion was done using the pre-trained ‘english-g2p’ model available within the Montreal Forced Aligner. The pronunciation lexicon, transcriptions, and audio file were aligned using the pre-trained ‘english’ acoustic model. The resulting annotations were visually examined in PRAAT (Boersma and Weenink, 2021) and manually adjusted when necessary. Phoneme onsets and word onsets predictors were modeled as impulses at the onset of each phoneme and word, respectively.

#### Phoneme surprisal and cohort entropy

Phoneme surprisal and cohort entropy reflect information-theoretic properties of the phoneme sequence in its lexical context, and are widely used in neural word processing analysis (Brodbeck et al., 2018; Gillis et al., 2021; Gwilliams et al., 2022). Phoneme surprisal quantifies the level of probabilistic surprisal associated with the current phoneme, given its occurrence after the sequence of phonemes preceding it within the current word. On the other hand, cohort entropy captures the level of uncertainty of remaining lexical candidates that match the observed phoneme sequence. Mathematically, phoneme surprisal for a given position *i* within a word is defined as the 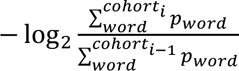 and cohort entropy is defined as 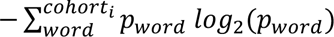. Here, *cohort_i_* refers to the set of words that are compatible with the phoneme sequence from the beginning of the word to the *i^th^* phoneme, and *p_word_* is the probability of the word derived from the *wordfreq* Python library (Speer et al., 2018). The *wordfreq* python library is based on the Exquisite Corpus (https://github.com/LuminosoInsight/exquisite-corpus) and covers a broad range of words that appear at least once per 100 million words. The phonetic lexicon for each word was extracted from the CMU pronouncing dictionary, available at http://www.speech.cs.cmu.edu/cgi-bin/cmudict. The final corpus used comprised of all the words that were included in both the CMU dictionary and the *wordfreq.* Cohort entropy and phoneme surprisal values were computed for each phoneme and represented as impulses at phoneme onset, scaled by the corresponding value. These two predictors were similar in the non-speech, scrambled, and narrative passages as they included meaningful words. However, they showed different distributions between meaningful words and non-words as illustrated in Figure 1-2B. As expected, phoneme surprisal exhibited a greater proportion of highly surprising phonemes for non-words, whereas cohort entropy displayed more zeros for non-words, since the potentially available lexicon usually becomes empty after some number of phonemes.

#### Unigram word surprisal and contextual word surprisal

Analogous to the phoneme level surprisal predictor, two different measures of word level surprisal were estimated: word frequency (quantified as unigram word surprisal) and contextual word surprisal. Unigram word surprisal measures how surprising a word is independent of the context and is based on the probability distribution of individual words computed from word frequencies (*wordfreq)*. Unigram word surprisal for each word is calculated by − log_2_(*p_word_*) and represented as an impulse at each word onset, scaled by the unigram word surprisal value (hereafter: word frequency). In contrast, contextual word surprisal depends on the preceding context and reflects how surprising the current word is given the previous context. Contextual word surprisal was estimated using the open source, pre-trained, and transformer-based (Vaswani et al., 2017) large language model GPT-2, implemented in the Hugging Face environment (Wolf et al., 2020). Each 1-minute-long passage was preprocessed (removing punctuation and converting to lower case, with the exception of proper nouns), tokenized using byte-pair encoding (Sennrich et al., 2016), and provided to the neural network model. The tokens could represent either complete words or sub-words. The final layer of the model was utilized to calculate the word surprisal. This final layer outputs prediction scores for each token in the vocabulary, indicating the likelihood of it being the next word given the preceding tokens (context) that extends all previous tokens, extending to a theoretical maximum of 1024 tokens (though the maximum number of words in the passages here was less than 220). The prediction scores were subjected to a SoftMax transformation to compute probabilities. The current word probability was determined by the probability associated with its corresponding token. In cases where words span over multiple tokens, word probability was computed by the joint probability of those tokens. Contextual word surprisal was computed as – log_2_(*P_word_*|*context*) and represented as an impulse at each word onset, scaled by the corresponding contextual word surprisal of that word. The word frequency and contextual word surprisal values were calculated only for the scrambled and narrative passages since they were not defined for non-words. However, as can be seen from Figure 1-2C, a high correlation between contextual word surprisal and word frequency was observed for the scrambled word condition (*r*(741) = 0.91, *p* < 0.001), suggesting that, as would be expected, contextual word surprisal collapses to word frequency when the context fails to provide informative cues for predicting the next word. Due to this very strong correlation between these two predictors in the scrambled passages, the contextual word surprisal predictor was excluded from the TRF modelling there and only the more conservative word frequency was used.

### Forward model (Temporal Response Functions)

The forward model approach referred to as temporal response function analysis (Lalor et al., 2009) was used to estimate how a set of predictor variables relates to the source localized MEG data. The model for each neural source is defined as:

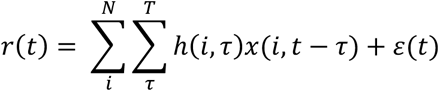

Where *r*(*t*) is the neural response at time *t*, *x*(*i*, *t*) is the *i*^th^ predictor time series, and *ε*(*t*) is the residual neural response not explained by the model. The TRF, ℎ(*i*, *τ*), is a filter that describes the linear relationship between the predictor time series and neural source time series (input and output) at different time lags within the integration window [*τ*, *T*]. In this model, each time lag of each predictor competes against each other to explain variance of the neural response, which results in larger TRF model weights associated with greater contributions to the explained variance. The TRF model weights were estimated by minimizing the mean absolute difference between actual (*r*(*t*)) and predicted 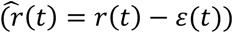 neural response.

To compute TRFs for each subject, condition, and at each source dipole, the eight trials per condition (total 8 minutes) were concatenated and the boosting algorithm (David et al., 2007) was employed. Prior to boosting, L1 standardization was performed on both the predictors and neural responses by subtracting the mean and dividing by the mean absolute value. TRF lags from −20 ms to 800 ms were used, with a basis of 50 ms Hamming windows employed to smooth the otherwise overly sparse TRFs. TRF estimation used four-fold cross-validation, where two folds were allocated for training, one-fold for validation and one-fold for testing. For each testing fold, each of the remaining three partitions served as a validation set, resulting in three TRFs per testing fold. These three TRFs were averaged to generate one average TRF per testing fold, which was then used to compute the prediction accuracy against the testing set. The TRFs from all testing folds were averaged to generate a single TRF for per source dipole. The predicted responses from each testing fold were concatenated to calculate a single prediction accuracy for each source dipole.

### Phonetic feature modelling

Before starting, we first analyzed how the phonetic features, phoneme onset, phoneme surprisal and cohort entropy should best be modeled, since different previous studies have used different approaches: modeling word-initial phonemes as separate features (Brodbeck et al., 2018); including word-initial phonemes with all other phonemes in phoneme surprisal and cohort entropy (Gaston et al., 2023); and including word-initial phoneme only in phoneme onset (Gillis et al., 2021). We compared models with and without word-initial phoneme onset on a base model with envelope spectrogram, envelope onset and word onset. The model with the word-initial phoneme onset showed better prediction accuracy compared to a model without the word-initial phoneme onset (*t*_*max*_ = 5.31, *p* < 0.001). To test for the phoneme surprisal and cohort entropy, we compared the three models by including, excluding, or separately modelling the word-initial phoneme, using a base model with gammatone envelope spectrogram, onset spectrogram and word onset. Model comparisons with adjusted r-squared revealed that including the word-initial phoneme yield the best prediction accuracies for both phoneme surprisal (1 *vs* 2 ∶ *t*_*max*_ = 4.38, *p* < 0.001, 1 *vs* 3 ∶ *t*_*max*_ = 3.81, *p* = 0.02) and cohort entropy (1 *vs* 2 ∶ *t*_*max*_ = 5.07, *p* < 0.001, 1 *vs* 3 ∶ *t*_*max*_ = 4.78, *p* = 0.02). We therefore opted to include the word-initial phoneme in the phonetic feature modelling.

### TRF peak extraction

TRFs showed prominent peaks with a distinct polarity at distinct latencies, reflecting major processing stages along the speech and language processing pathway. The amplitudes and latencies of these peaks served as the strength of neural processing at the corresponding stage. To investigate how neural auditory processing stages differ based on the linguistic content of the stimuli, the peak amplitudes and latencies were compared across passage types.

First, we identified the time windows for the main peaks associated with each predictor and their respective polarities based on a combination of prior literature and visual inspection of the group averaged TRFs (Brodbeck et al., 2018; Gillis et al., 2021; Keshishian et al., 2023). The time windows for each predictor were 1) Envelope: Early (20-130 ms), Late (70-180 ms); 2) Envelope onset: Early (20-170 ms), Late (70-240 ms); 3) Phoneme onset: Early (40-200 ms), Late (120-410 ms); 4) Phoneme surprisal: Early (40-200 ms), Late (110-470 ms); 5) Cohort entropy: Early (40-120 ms), Middle (140-350 ms), Late (260-600 ms); 6) Word onset: Early (40-200 ms), Middle (220-350 ms), Late (310-650 ms); 7) Word frequency: Early (40-300 ms), Late (310-610 ms); 8) Contextual word surprisal: Early (40-300 ms), Late (310-610 ms). Early and middle peaks have positive current polarity while the late peak is a negative current polarity peak (respectively, directed out of, or into, upper surface of the superior temporal gyrus).

A peak-picking algorithm was developed to pick the maximum peaks with the corresponding polarity within the given time window. The algorithm followed these steps: 1) TRFs were aggregated across the source ROIs by taking the absolute sum; 2) Peaks within the given time window were identified; 3) Selection of the maximum peak that aligns with the target polarity by checking the source current polarity relative to cortical surface in the transverse temporal region in the original source TRFs; 4) If none of the peaks satisfied the polarity constraint, the minimum of the average TRF in the given time window was used as the peak amplitude, and the latency was set to NaN (not a number). A small number of peaks (<1.5 %) were further manually adjusted where appropriate.

### Statistical analysis

Statistical analysis was performed in R (R Core Team, 2020) version 4.0 and Eelbrain. The significance level was set at *α* = 0.05.

Significance of each speech feature over and beyond other features was evaluated by comparing full and reduced models. The full models for modulated noise and non-words included: gammatone envelope, envelope onset, phoneme onset, phoneme surprisal, cohort entropy and word onset. Additionally, the scrambled passages also included word frequency; the narrative passages included both word frequency and context-based word surprisal. Each reduced model included all the features of the full model, except excluding the single predictor under investigation. The proportion of explained variance between the full and reduced model at each current source dipole were tested using mass-univariate one-tailed paired sample t-test with threshold-free cluster enhancement (TFCE (Smith and Nichols, 2009)) with a null distribution based on 10,000 permutations of model labels.

Hemispheric lateralization of each feature was performed to examine the lateralization of each speech feature processing. The explained variance maps for each feature were transformed to a common space by first morphing to a symmetric brain template ‘fsaverage_sym’ and consecutively morphing the right hemisphere to the left hemisphere. The explained variance between left and right hemispheres were tested using mass-univariate two-tailed paired sample t-tests with TFCE.

TRF amplitude comparisons were performed using repeated measures ANOVAs and using post hoc paired sample t-tests with correction for multiple comparisons using false discovery rate correction. To ensure unbiased TRF comparison across passage types, TRFs were generated from a similar number of predictors across passage types. The effect sizes for paired sample t-tests were calculated using Cohen’s d (*d*) (*d* = 0.2 indicates a small effect, *d* = 0.5 indicates a moderate effect, and *d* = 0.8 indicates a large effect).

Statistical summary tables are reported in Extended data.

### Data availability

The raw MEG data, stimulus materials, analysis codes, intermediate results, and behavioral responses are available to download through reviewer sharing link, https://datadryad.org/stash/share/EJjwUNsN9k3ToO68sI1yAkjxR3Izfpyj00lLPNSvTic. Code and dataset supporting the findings of this paper will be shared once the paper is accepted.

## Results

Using acoustic stimuli with similar prosody and rhythm but progressing from lacking any linguistic information (speech modulated noise) to possessing well-formed phonemes but no more (non-words), to possessing well-formed words but no larger scale context (scrambled), to fully well-formed linguistic information (narrative), we trace changes in the neural response dynamics as speech and speechlike sounds are eventually turned into language with full meaning in an ecologically valid setting. We first present the emergence of features, from acoustic to sentence-level, as speech processing unfolds incrementally. Next, we show how the hemispheric lateralization progresses from acoustic to sentential processing. Finally, we explore the temporal dynamics of speech processing using temporal response function profiles.

### Emergent features of speech processing

The present study first aimed to investigate the emergence of neural speech processing in response to varying levels of speech and linguistic information in the sensory input, by testing which speech representations are tracked by the brain response for each passage type. The test for significance of each speech representation (predictor) was done by comparing explained variance within pairs of models, one with all predictors included and the other for which the test predictor (speech representation of interest) was excluded; the test predictor was denoted as significant if the difference in explained variance was statistically significant. The full model employed for passages using speech-modulated noise and non-words included predictors for: gammatone envelope spectrogram, gammatone onset spectrogram, phoneme onset, word onset, phoneme surprisal, and cohort entropy. The model for scrambled word passages additionally included word frequency, and for narrative passages additionally incorporated both word frequency and contextual word surprisal. For the non-word passages, neither word frequency nor contextual word surprisal could be applied as there were no real words. The studies using comprehensible and incomprehensible language (Gillis et al., 2023; Tezcan et al., 2023) have shown that higher level word features (word frequency, word entropy, and contextual word surprisal) are not encoded for incomprehensible language, the explicit quantification of differences between non-words and meaningful words was not conducted in our models. This is due to the unavailability of word frequency for non-words. Any word frequency defined for non-words would be uniform across all non-words, and therefore identical to the word onset predictor already included. In the scrambled word passages, where context does not provide meaningful cues, contextual word surprisal collapsed to the word frequency (see Methods, predictor variables); therefore, only the word frequency was used, since the explained variance by contextual word surprisal in the absence of coherent meaning is more conservatively ascribed to that of word frequency. Statistical summary tables are reported in Figure 2-1.

Model comparison results for all passage types are illustrated in Figure 2. In the modulated noise condition, only the acoustic features, specifically the gammatone envelope spectrogram (*t*_*max*_ = 6.92, *p* < 0.001) and gammatone onset (*t*_*max*_ = 5.79, *p* < 0.001), contributed significantly to the observed neural data variance explained, i.e., significantly improving the full model fit over the reduced model. Conversely, none of the linguistic predictors, phoneme onset (*t*_*max*_ = 3.30, *p* = 0.07), word onset (*t*_*max*_ = 2.46, *p* = 0.91), phoneme surprisal (*t*_*max*_ = 1.51, *p* = 1.0), and cohort entropy(*t*_*max*_ = 1.82, *p* = 0.99) showed a significant contribution to the model’s predictive power. However, in the presence of low-content speech stimuli, whether non-words or scrambled words, in addition to these acoustic features, linguistic segmentation responses (phoneme and word onset) and statistically based linguistic features (phoneme surprisal and cohort entropy) also significantly contributed to the model’s predictive power (non-words: gammatone envelope (*t*_*max*_ = 11.90, *p* < 0.001), gammatone onset (*t*_*max*_ = 9.37, *p* < 0.001), phoneme onset (*t*_*max*_ = 7.25, *p* < 0.001), phoneme surprisal (*t*_*max*_ = 5.60, *p* < 0.001), cohort entropy (*t*_*max*_ = 6.83, *p* < 0.001), word onset (*t*_*max*_ = 6.90, *p* < 0.001); scrambled words: gammatone envelope (*t*_*max*_ = 10.97, *p* < 0.001), gammatone onset (*t*_*max*_ = 10.68, *p* < 0.001), phoneme onset (*t*_*max*_ = 6.43, *p* < 0.001), phoneme surprisal (*t*_*max*_ = 7.13, *p* < 0.001), cohort entropy (*t*_*max*_ = 8.60, *p* < 0.001), word onset (*t*_*max*_ = 6.17, *p* < 0.001)). These results indicate that the acoustic features represented by the gammatone envelope and onset spectrograms are encoded in the brain regardless of the intelligibility of the sensory input, whereas linguistic features are tracked by the brain as soon as the linguistic units or linguistic unit boundaries are intelligible, regardless of any higher-level meaning.

**Figure 2.**
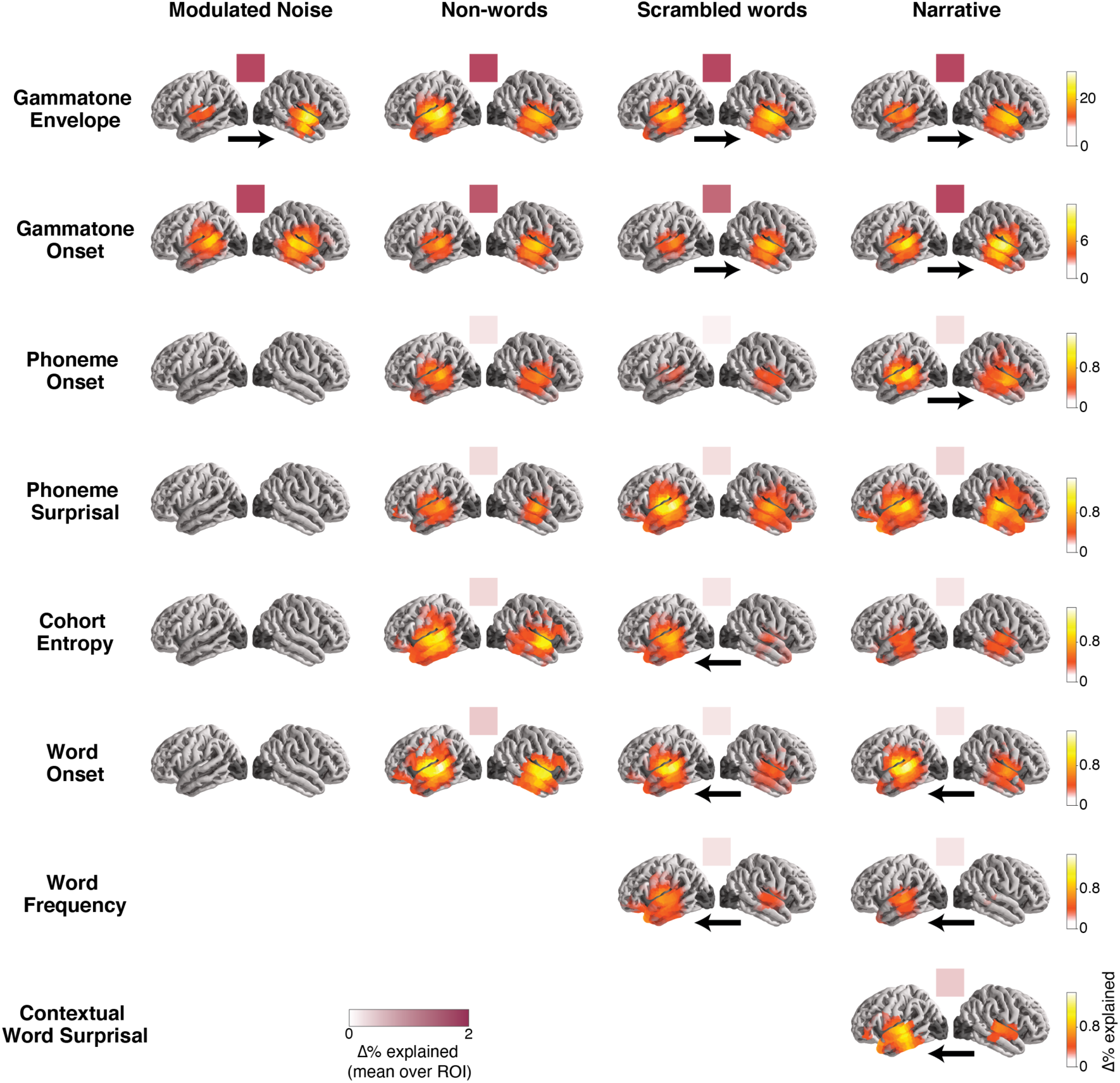
Emergence of hierarchical speech processing. Anatomical brain plots visualize the cortical regions where each respective predictor significantly contributes to the model fit. Colored squares above the anatomical plots indicate average explained variance over frontal, temporal, and parietal regions. Black arrows below anatomical plots indicate significant hemispheric asymmetry. The first two rows show that acoustic features are represented in the brain irrespective of the passage type and intelligibility. Later rows show that linguistic features are tracked only when the linguistic feature boundaries are intelligible, irrespective of any higher-level (e.g., sentential meaning). When the context supports higher-level meaning above and beyond that of individual words, contextual word surprisal is additionally represented in the brain. Lower-level feature processing is more right-lateralized, while higher level feature processing is more left-lateralized.

Furthermore, model comparisons conducted on both scrambled (*t*_*max*_ = 6.67, *p* < 0.001) and narrative (*t*_*max*_ = 6.48, *p* < 0.001) passages revealed that when the words are individually meaningful, and irrespective of the structured coherence of the passages, the brain significantly tracked word frequency (absent of context). This suggests that the brain is sensitive to both the overall predictability and integration of individual words, regardless of the overall coherence of the passage. Additionally, for narrative passages, where structured contextual meaning was present, the brain exhibited substantial additional tracking of contextual word surprisal (*t*_*max*_ = 5.48, *p* < 0.001), over and beyond word frequency. This context-based word surprisal processing represents a higher-level processing that involves integration of linguistic and syntactic information to construct a structured meaning (Heilbron et al., 2022; Caucheteux et al., 2023). Model comparison between word frequency and contextual word surprisal in narrative passages additionally verified that contextual word surprisal is better encoded in the brain than word frequency (*t*_*max*_ = 4.70, *p* = 0.02). These results indicate that the brain maintains both context-free and contextual representations during speech understanding (Brodbeck et al., 2022), but contextual-level information is more strongly represented.

The anatomical distribution of the neural sources processing this hierarchy of speech processing was observed in locations consistent with an origin in Heschl’s gyrus (HG), spreading to the superior temporal gyrus (STG) and much of temporal lobe (Figure 2). For higher-level linguistic features including phoneme surprisal, cohort entropy, word onset, word frequency, and contextual word surprisal, the feature representations additionally extended to left frontal regions.

### Lateralization of speech feature processing

Hemispheric lateralization of auditory and speech processing has been widely studied and is of great interest, but results still show much variability across different studies (Peelle, 2012). Therefore, we also examined the lateralization of neural speech feature processing for each passage type and speech feature. Instances of statistically significant lateralization are indicated by arrows in Figure 2. Lateralization varied depending on the passage type and specific speech feature. Overall, lower-level speech feature processing exhibited a bilateral and right lateralized pattern (narrative: gammatone envelope (*t*_*max*_ = −5.04, *p* < 0.001), gammatone onset (*t*_*max*_ = −4.36, *p* = 0.02), phoneme onset (*t*_*max*_ = −4.57, *p* = 0.005)) in the sources spanning in most of the temporal lobe, whereas higher-level speech feature processing were more left lateralized (narrative: word onset (*t*_*max*_ = 3.21, *p* = 0.02), word frequency (*t*_*max*_ = 3.23, *p* = 0.03), contextual surprisal (*t*_*max*_ = 3.30, *p* = 0.02)) in superior temporal gyrus (STG), anterior temporal lobe and extending into frontal cortex. On the other hand, phoneme-level feature processing displayed a more bilateral pattern (narrative: phoneme surprisal (*t*_*max*_ = −2.01, *p* = 0.82), cohort entropy (*t*_*max*_ = 2.38, *p* = 0.63)). These results suggest distinct specialization of hemispheric regions for the processing of lower-level acoustic information vs. higher-level linguistic analysis.

Interestingly, the non-word passages showed predominantly bilateral responses across the different speech features (gammatone envelope (*t*_*max*_ = 4.33, *p* = 0.06), gammatone onset (*t*_*max*_ = −4.17, *p* = 0.04), phoneme onset (*t*_*max*_ = 3.58, *p* = 0.08), phoneme surprisal (*t*_*max*_ = 2.72, *p* = 0.29), cohort entropy (*t*_*max*_ = 3.92, *p* = 0.06), word onset (*t*_*max*_ = 2.58, *p* = 0.39)), suggesting a more symmetrical hemispheric engagement of neural resources in non-word processing.

### Effect of context on progression of neural speech processes: early and late

Neural responses obtained using MEG, with its fine-grained time resolution, often provide even greater insight from the temporal progression of cascading neural processes than from their anatomical locations.

Having tested which types of speech-feature processing occurs in different contexts and in different anatomical regions, we then investigated how these contextual factors also influence the underlying neural mechanisms, associated with the processing of each speech feature, in the time domain. To this end, we utilized TRF analysis that describes how the brain responds to each predictor over a range of latencies. To compare the TRFs between passage types, TRF magnitudes over the brain sources were aggregated. Analogous to ERP responses to punctate sounds, that exhibit distinct peaks at specific latencies characterized by their current polarity, so also do these TRFs, representing the direction and strength of the neural current response to each predictor, at various latencies. The dominant TRF peaks were identified and compared across passage types using repeated measures ANOVA (post hoc paired sample t-tests corrected for multiple comparisons using the false discovery rate method). To ensure unbiased TRF comparison across passage types, TRFs were generated from the same number of predictors. Peak latencies were also compared (Figure 6A), and unless otherwise mentioned, no significant differences were found for latencies. Figures 3, 4 and 5 illustrate average TRFs and their main peaks, and the accompanying bar plots provide a comprehensive comparison across the different speech passage types. The results presented in these figures show either only left or right hemisphere responses, so as not to overwhelm the figures; however, full analysis results are included in the extended data (Figure 3-(1-3), Figure 4-(1-6), Figure 5-(1-3)).

**Figure 3.**
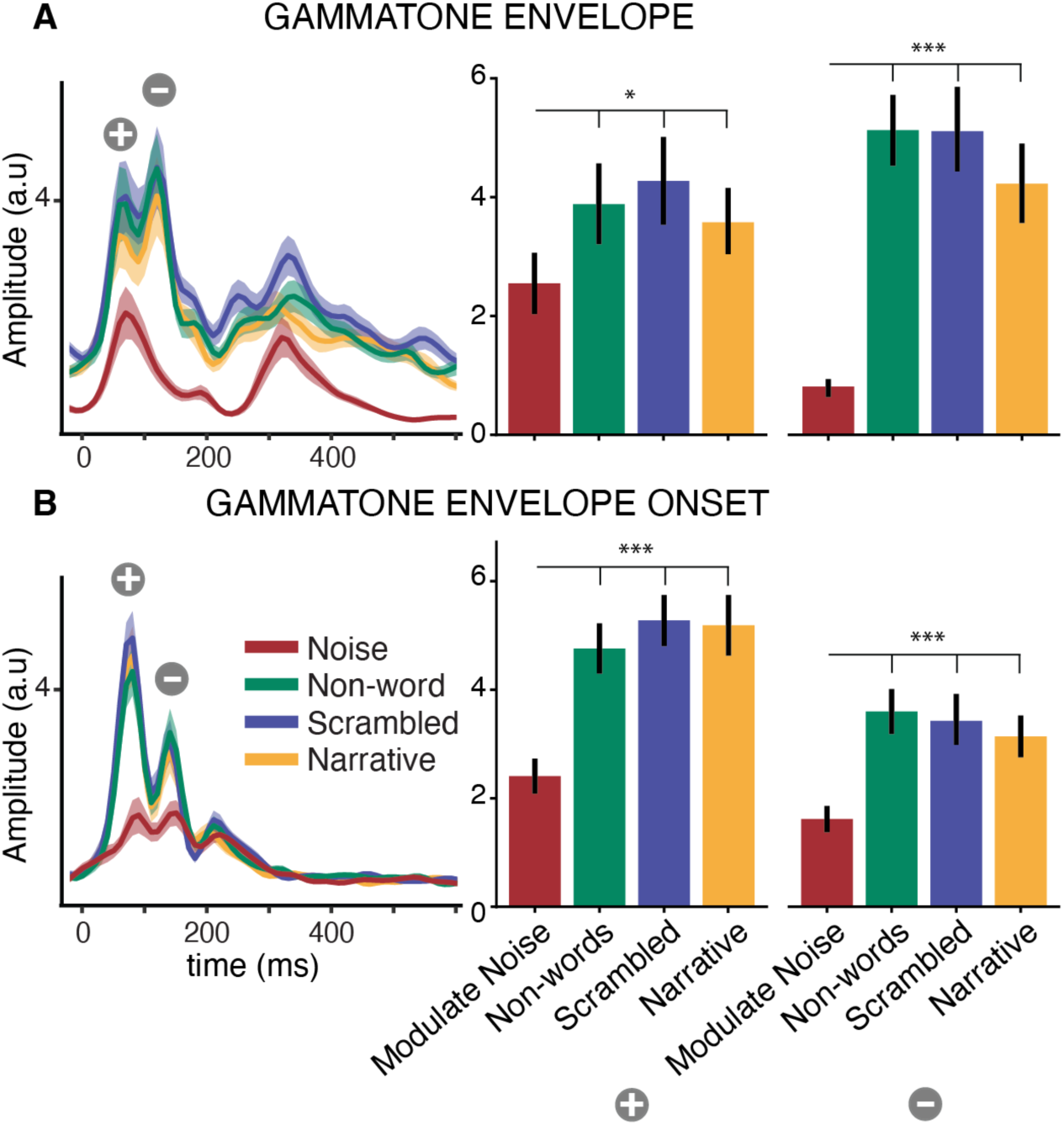
Neural responses to acoustic features. (A). Gammatone envelope and (B). gammatone envelope onset responses. Left panels show the TRF magnitude aggregated over sources and subjects, by passage type. The TRFs exhibit an early positive and a late negative polarity peak indicated by 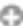 and 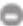 respectively. The right panel bar plots (mean±standard error (SE)) compare the peak amplitudes, first early then late, across passage types. Both early and late responses are stronger for speech compared to non-speech (noise). Only right hemisphere results shown (see Figure 3-1 for both hemispheres and individual data points). *p<0.05, **p<0.01, ***p<0.001

**Figure 4.**
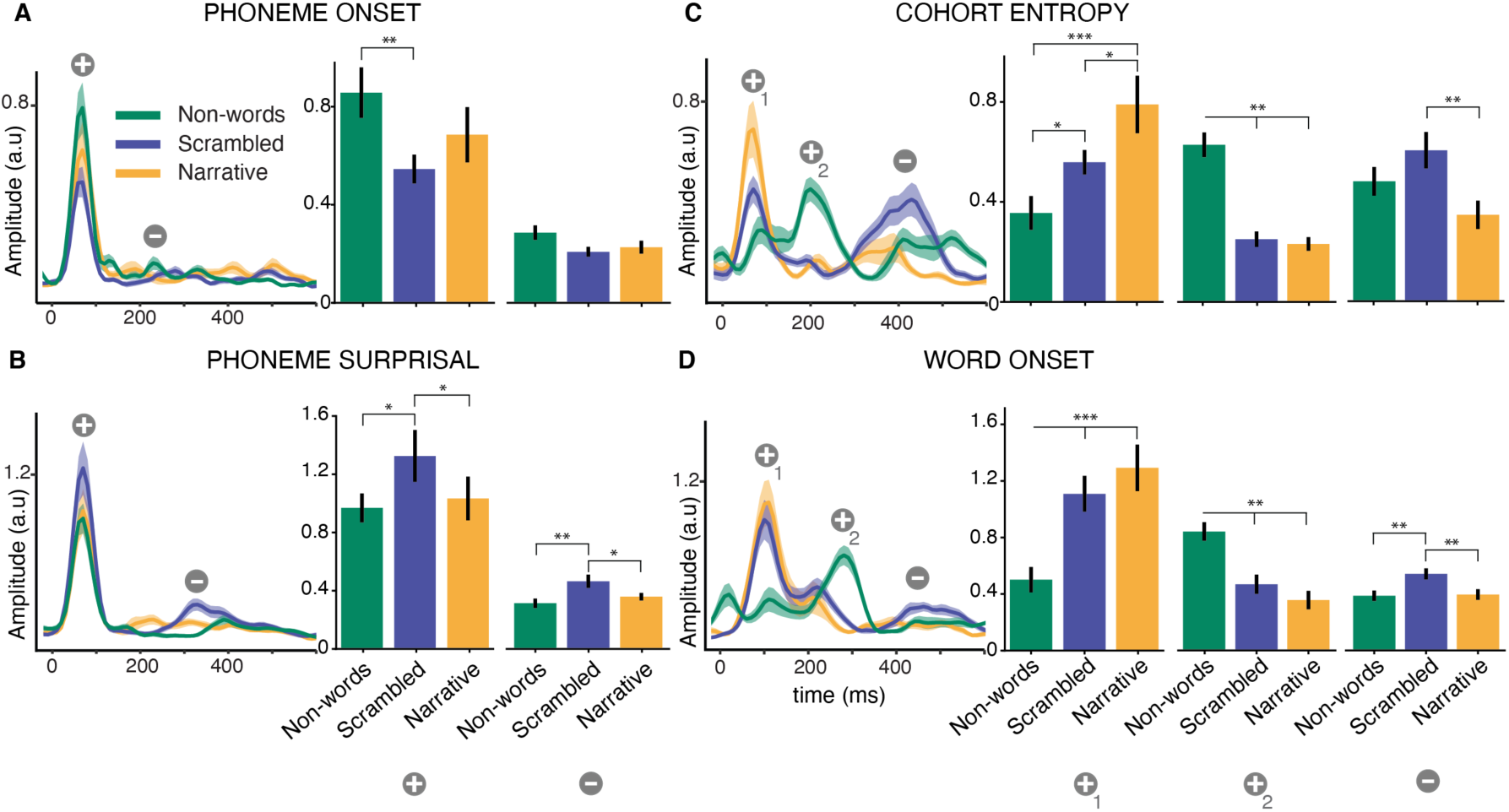
Neural responses to sub-lexical and word onset speech features. (A). Phoneme onset, (B). phoneme surprisal, (C). cohort entropy, and (D) word onset (TRF magnitude plots and TRF peak bar plots as in Figure 3). TRFs exhibit an early positive and a late negative polarity peak indicated by 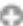 and 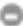 respectively. For both word onset and cohort entropy responses, non-words showed a robust positive polarity peak between early and late peaks. These early, middle, and late peaks are indicated by 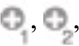 and 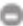 respectively. The bar plots compare the peak amplitudes across passage types. Only left hemisphere results are shown here (see Figure 4-1 for both hemispheres and individual data points). Overall, the early responses were very differently modulated by the linguistic content. The middle peak (second positive polarity peak) was strongest for non-words, while the late peak (negative polarity) was strongest for scrambled passages.

**Figure 5.**
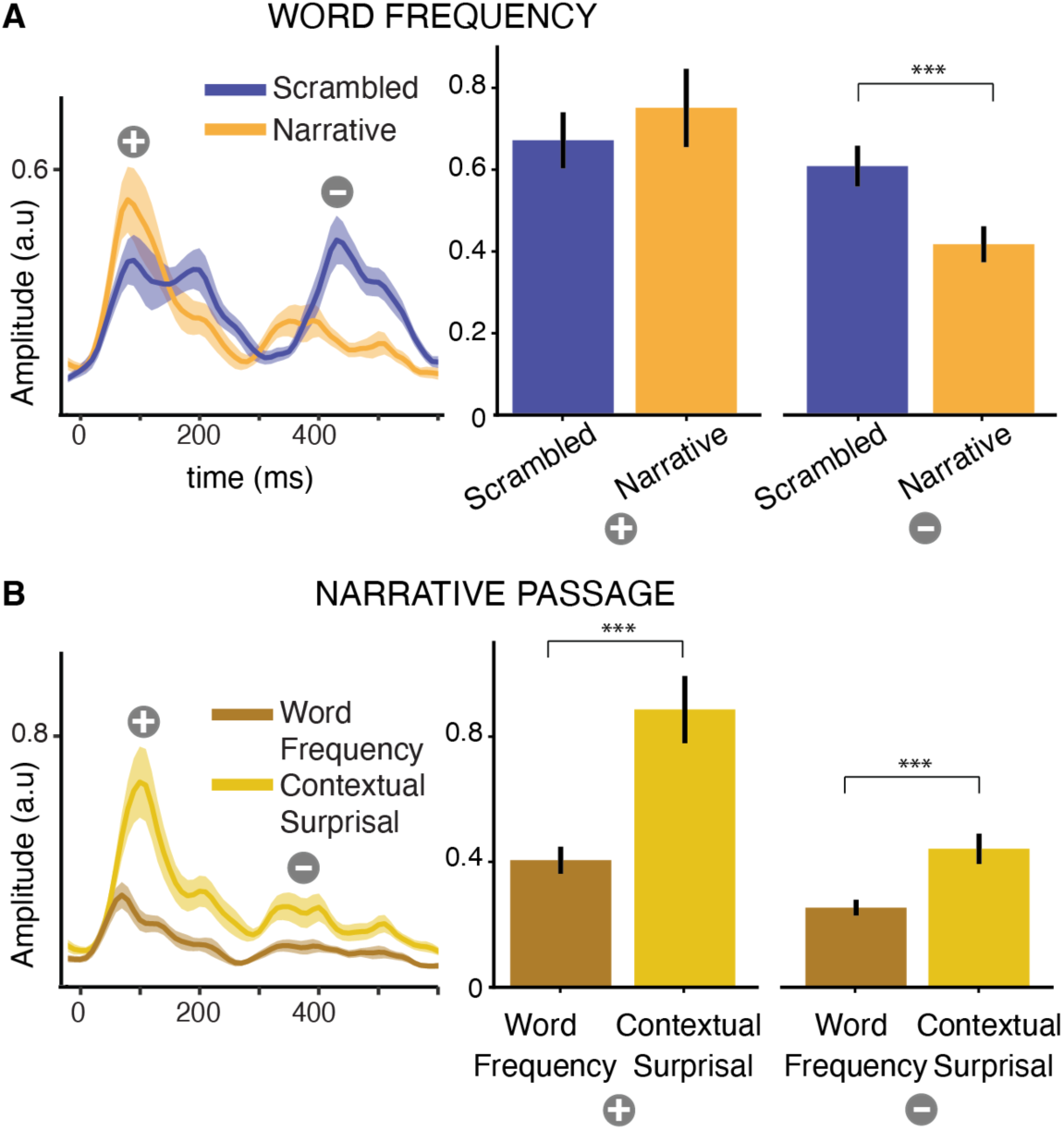
Neural responses to lexico-semantic features. (A). Word frequency (B). Word frequency and contextual word surprisal for the narrative passage (TRF magnitude plots and TRF peak bar plots as in Figure 3). The TRFs exhibit an early positive and a late negative polarity peak indicated by 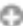 and 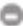 respectively. Only left hemisphere results are shown here (see Figure 5-1 for both hemispheres and individual data points). The late word frequency responses (N400-like) are stronger for scrambled passages compared to narrative passage. Contextual word surprisal responses are stronger compared to word frequency responses. Note that the peak amplitudes for word frequency in (A) and (B) are different, as the TRF model in (A) does not include a separate predictor for contextual surprisal.

**Figure 6.**
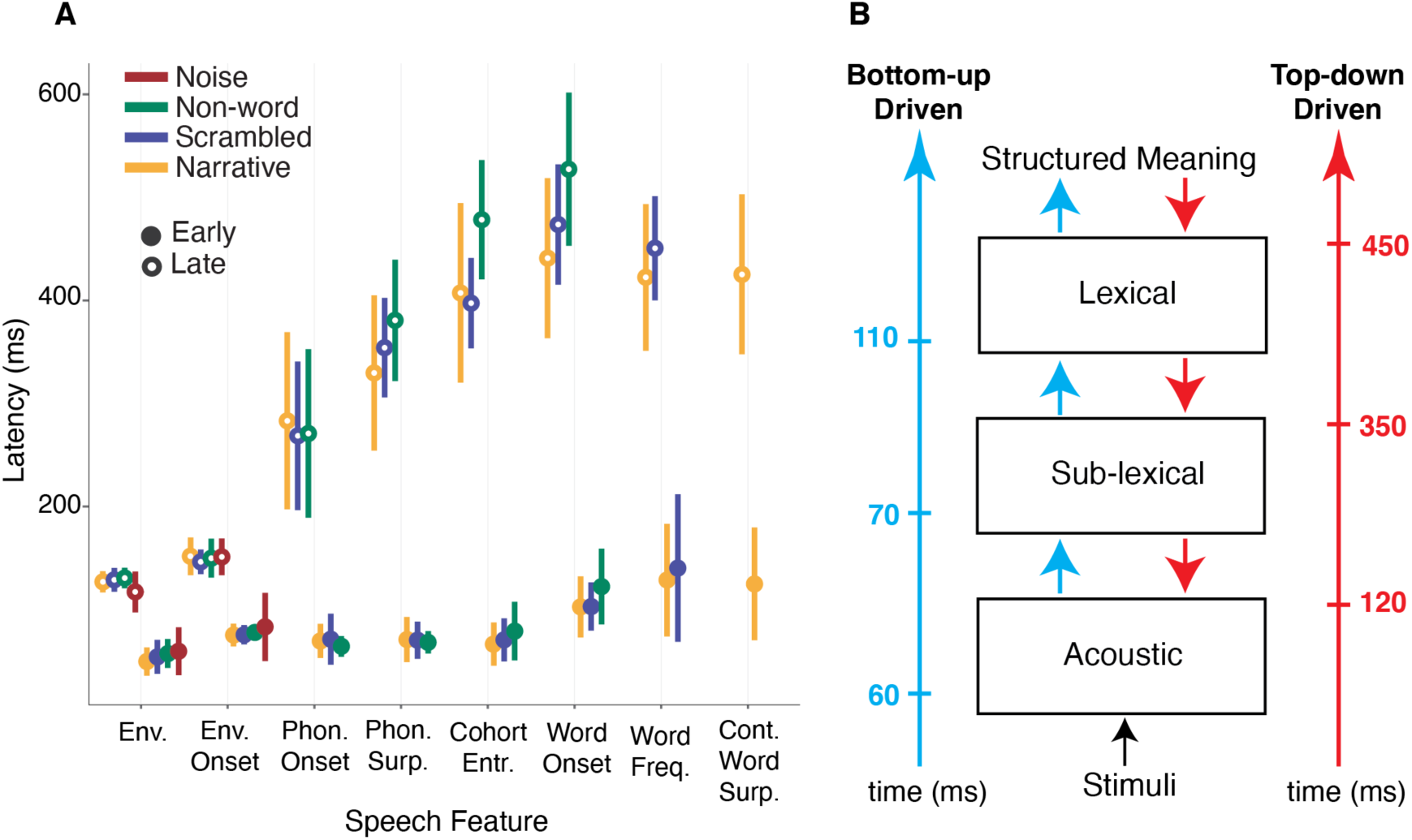
Temporal profile of speech feature processing. (A). Latency of both early and late processing stages associated with each feature processing. As the features go up in hierarchical acoustic and linguistic structures both early and late peak processing show longer latencies, with all measures obtained simultaneously from the same continuous stimuli. (B). Schematic summary of the bottom-up and top-down temporal profiles associated with earliest bottom-up and top-down mechanisms at each level as outlined in the final section of the discussion. Acoustic = [Envelope, Envelope Onset], Sub-lexical = [Phoneme Onset, Phoneme Surprisal], Lexical = [Cohort Entropy, Word Onset, Word frequency, Contextual Word Surprisal]. Bottom-up mechanisms correspond to early processing stages, while top-down mechanisms emerge in the late processing stages, as inferred from their timing and modulation by linguistic content. Latencies marked are derived from envelope responses, phoneme surprisal responses, and word onset responses for each level.

Neural responses to acoustic features (Figure 3) showed two prominent peaks: an early peak with a positive current polarity, and a late peak with a negative current polarity. These two peak latencies for the gammatone envelope were ∼60 ms and ∼120 ms respectively, while for the gammatone onset feature, peak latencies were ∼70 ms and ∼150 ms (c.f. the early (P1) and late (N1) peaks of an auditory ERP). The late responses showed a predominantly right hemispheric lateralization (*p* < 0.001). When comparing these two neural responses across passage types, we found that neural responses to speech passages were stronger compared to the non-speech modulated-noise (*p* < 0.001). This effect was smaller for the right hemisphere early responses (left: early: *d* = 1.06, late: *d* = 1.120; right: early: *d* = 0.47, late: *d* = 1.20). The TRF amplitude differences between speech and non-speech passages, even at baseline, could be driven by the differences in statistics of the predictor variables (see Figure 1-2), and the engagement of more cortical areas for speech processing, as shown in Figure 2. It was also observed that the late peak was nearly absent in the modulated noise responses. When comparing the envelope onset responses among the speech passages, no significant differences were observed (*p* > 0.2). However, for envelope responses significant differences were found across speech passages in the left hemisphere. Early responses were smaller in narrative passages compared to scrambled and non-words (*p* < 0.001), whereas late responses were stronger in non-words compared to meaningful words (*p* < 0.02). Even though envelope and envelope onsets are temporally related, these stark differences observed in between them in response to passage type suggest that envelope and envelope onset tracking arise from quite different neural mechanisms (Hamilton et al., 2018).

The analysis of phoneme onset responses (Figure 4A) also revealed a robust early positive polarity peak with ∼70 ms latency; the substantially later peak at ∼250 ms latency was noisy and not robust across subjects (Di Liberto et al., 2015). When comparing the peak amplitudes across passage types, no significant differences were observed in the right hemisphere for late responses. In the left hemisphere, early responses were stronger for non-words compared to scrambled passages (*p* = 0.002).

Phoneme surprisal (Figure 4B) also showed two prominent peaks: an early positive polarity peak at ∼70 ms and a late negative polarity peak at ∼350 ms. Similar to phoneme onset responses, significant differences between passage types were found only in the left hemisphere. Both the early (*p* = 0.03) and late (*p* < 0.03) peaks were stronger in response to scrambled words compared to narrative and non-word passages.

For cohort entropy responses (Figure 4C), two main processing mechanisms were observed for the scrambled and narrative passages: an early positive peak at ∼70 ms and a late negative peak at ∼380 ms. However, non-word passages showed a robust intermediate positive polarity peak at ∼200 ms. Therefore, three peaks were identified as early, middle and late responses. The early peak was stronger for non-words compared to scrambled (*p* = 0.01) and to narrative (*p* = 0.02), while the middle peak was stronger in non-words compared to meaningful words (*p* < 0.001). In contrast, the late peak was stronger in scrambled words compared to narrative (*p* = 0.009); additionally, this peak was delayed for non-words compared to meaningful words (*p* < 0.001). Finally, the early cohort entropy responses were left lateralized for meaningful words (*p* = 0.002), middle non-word responses (*p* = 0.03) and late scrambled word responses (*p* = 0.001).

Analogous to cohort entropy responses, word onset responses (Figure 4D) displayed two main peaks for both scrambled and narrative passages, while a middle peak was evident for non-words. Both early and middle peaks, occurring at ∼100 ms and at ∼200 ms respectively, exhibited a positive polarity. In contrast, the broad late peak at ∼450 ms showed a negative polarity, resembling a characteristic N400 response. The early peak was stronger for meaningful words compared to non-words (*p* < 0.001), whereas this effect was reversed for the middle peak (*p* < 0.001). Interestingly, no significant differences were observed between the scrambled and narrative passages for both early (*p* = 0.09) and middle (*p* = 0.07) peaks. Remarkably, the late peak exhibited greater strength in response to scrambled words compared to non-words and narrative passages (*p* = 0.003). Moreover, the late peak latency was significantly delayed in the progression from narrative to scrambled (by ∼30 ms, *p* = 0.02) to non-words (by ∼50 ms, *p* = 0.002). Additionally, consistent with the explained variance lateralization comparisons, the non-words early and middle responses showed bilateral response (*p* = 1.0), while in meaningful words, the early responses were left lateralized (*p* = 0.001).

In the above analysis, we conservatively separated the early and middle peaks in cohort entropy and word onset responses into different processing stages, due to the considerable temporal separation between them. However, because of the strong similarity between the peak amplitudes and polarity, we also performed a separate analysis where the positive peaks (early and middle) were grouped together. In this analysis no significant differences in peak amplitudes were observed across the passage types (cohort entropy: *p* > 0.08, word onset: *p* > 0.06); as expected, latency comparisons revealed that the peak is delayed in non-words compared to meaningful words (*p* < 0.001).

Word frequency TRFs (Figure 5A) showed two main peaks, comparable to the early and late peaks observed in the word onset responses. Consistent with the explained variance lateralization, both peaks showed left hemispheric dominance. When comparing the peak strength between the scrambled and narrative passages, no significant differences were found for the early peak (*p* = 0.16). However, interestingly, the late peak in the scrambled word passages TRF was stronger (*p* < 0.001) and delayed by ∼30 ms (*p* = 0.04) compared to narrative passages.

The TRFs between word frequency and contextual word surprisal within the narrative passage were also compared (Figure 5B). Both predictors represent word surprisal and exhibit a similar range of values, facilitating a direct comparison. Both TRFs showed similar peaks at comparable latencies and were left lateralized (*p* < 0.001). In contrast to the similarity in peak timing, contextual word surprisal showed stronger amplitudes for both early (*p* < 0.001) and late (*p* < 0.001) peaks in both hemispheres when compared to word frequency, indicating contextual information is more robustly tracked.

The current analysis does have its limitations. Specifically, more fine-grained stages within the speech and language processing hierarchy, such as syntactic-only processing and semantic-only processing, were not included (due to experimental constraints related to recording durations). Additionally, other speech features, including but not limited to morphemes, function words, and content words, were not incorporated into the analysis. Further, the stimuli across passage types were not tightly controlled for linguistic structures. Investigating such aspects would indeed be a valuable direction for future research.

In summary, our TRF analysis revealed that the brain processes the hierarchy of acoustic and linguistic structures (from acoustics to context-based features) in a progression of neural stages, and with a characteristic temporal dynamic associated with each feature processing. As we ascend the hierarchy (when speech features become more abstract and less directly related to the acoustics), processing of features shows longer latencies for both early and late mechanisms (Figure 6A), suggesting a graded computation of features, over time, in the cortex (Keshishian et al., 2023), starting as early as ∼50 ms and extending to ∼500 ms. These mechanisms accumulate sounds features, analyze for lexical-semantic information, and integrate with the semantic context. Acoustic feature responses to speech were stronger compared to non-speech. Notably, the early and late peaks exhibited different modulations by linguistic content, consistent with representing different neural mechanisms. Even though different patterns were observed for early stage between passage types, the later stage trends were consistent for both phoneme level and lexical level, suggesting the late stage may represent similar neural mechanisms. Furthermore, the TRFs showed quite different peak latencies for non-words compared to meaningful words. For linguistic level features, the late TRF peak amplitudes were stronger for scrambled words compared to non-words and narrative passages. Additionally, cohort entropy and word level late processing were delayed from narrative to scrambled to non-words. Peak lateralization analysis was consistent with explained variance lateralization analysis: lower-level feature processing was more right-lateralized, while higher level feature processing was more left-lateralized.

## Discussion

Using multiple stimulus types with varying linguistic content, our results provide neural evidence for the progression of different speech features, hemispheric lateralization, temporal dynamics and neural mechanisms associated with each level and how they are further modulated by linguistic content. Our results complement and extend fMRI studies (Binder, 2000; Xu et al., 2005) by leveraging the temporal dynamics of feature processing and electrophysiological studies by investigating effects of linguistic content on neural tracking measures (Gillis et al., 2021).

We first showed that the brain separately represents hierarchical speech and linguistic structures, with emergence of these features from acoustics to contextual processing arising with the increasing contextual information necessary for language comprehension. Regardless of the stimulus type, acoustic envelope and envelope onsets are represented in the brain (Kubanek et al., 2013; Steinschneider et al., 2013; Oganian and Chang, 2019), reflecting a lower-level, initially bottom-up, sensory processing mechanism (Karunathilake et al., 2023). (Sub)-lexical features processes are activated as soon as (sub)-lexical units are recognizable and intelligible for linguistic process activation (Overath et al., 2015). Moving from non-words to meaningful words, results show the emergence of lexico-semantic processes, while avoiding the inherent confounds of instead using incomprehensible foreign languages (Gillis et al., 2023; Tezcan et al., 2023). Moving from scrambled words to narrative passages, our results also show the emergence of context-based word processing, robustly represented compared to non-contextual word predictions, indicating that the brain strongly incorporates context to predict the structured meaning in line with the predictive coding theories (Dambacher et al., 2006; Payne et al., 2015; Schrimpf et al., 2021). These two measures represent different cognitive operations, where context-based surprisal involves word retrieval based on contextual and syntactic information, whereas word frequency relies solely on sensory cues (Bentin et al., 1999; Huizeling et al., 2022).

Our lateralization results underscore the specialized contribution of each hemisphere to different levels of speech comprehension within a common stimulus and emphasize the brain’s flexibility in adapting to various linguistic and acoustic demands. The results demonstrate that pre-lexical auditory input analysis occurs in both hemispheres, with a right hemispheric advantage, and left lateralization emerges as lexical-semantic processing becomes involved (Overath and Paik, 2021). While lower-level acoustic processing has been identified as bilateral (Binder, 2000; Aiken and Picton, 2008), the right hemisphere’s extra involvement in acoustic level processing aligns with its specialization in acoustic analysis, including extraction of spectral and temporal features from auditory input (Ross et al., 1997; Poeppel, 2003; Ding and Simon, 2012a). The left lateralization for higher level responses is consistent with the well-established left hemisphere specialization for language functions (Hickok and Poeppel, 2004, 2007; Gow, 2012). Indeed, it is crucial to emphasize that numerous studies have reported different patterns of lateralization across task and language processes (Price, 2012; Fedorenko et al., 2012; Bradshaw et al., 2017). Interestingly, non-word processing exhibited bilateral responses at every level of processing, suggesting both hemispheres are engaged in the absence of successful lexical retrieval for non-word understanding (Bozic et al., 2010; Mai et al., 2016).

Critically, TRF analysis revealed multiple processing stages, with distinct temporal dynamics influenced by bottom-up and top-down driven mechanisms (Shuai and Gong, 2014; Arnal et al., 2016). These mechanisms at each stage are inferred from the timing and their modulation by linguistic complexity, as detailed in the following paragraphs, summarized in Figure 6B. Some predictors are inherently bottom-up, driven by sensory cues, while others reflect top-down influence shaped by linguistic experience (Gwilliams and Davis, 2022). For example, envelope responses, driven by sensory signal, are observed early for both speech and nonspeech stimuli, with late-stage processing emerging only for speech, suggesting top-down involvement of speech processing (Ding and Simon, 2012b; O’Sullivan et al., 2015). Similarly, linguistic level responses, which are not directly tied to the sensory signal, also show early and late responses. These responses arise from the listener’s internal language model, formed through lifelong linguistic exposure (Gwilliams and Davis, 2022). The early response points to a generalized feature regardless of the structure, reflecting the generation of predictions through a bottom-up driven mechanism driven by the statistics of the internal model. In contrast, prediction evaluation and adjustments occur during the late stage, resulting in modulation by linguistic content and highlighting the influence of top-down processing (Gwilliams and Marantz, 2015). It has been also seen that some predictive coding models may even be dominantly bottom-up, for example, in an auditory midbrain model of predictive processing (de Cheveigné, 2024).

Acoustic feature responses were stronger for speech compared to the non-speech (Binder, 2000; Nourski et al., 2019), attributable to the underlying acoustic differences (Karunathilake et al., 2023). Although it might appear from the current study that envelope onset responses are dominantly bottom-up, previous work has shown a strong top-down modulatory influence, especially from selective attention, on the later peak (Fiedler et al., 2019; Brodbeck et al., 2020).

Phoneme onset responses reflect more of a mixed acoustic-linguistic measure rather than a purely linguistic measure (Karunathilake et al., 2023). The early responses for non-words were enhanced for phoneme onset but were smaller for phoneme surprisal and cohort entropy compared to scrambled passages. It is also possible that differences in predictor distributions between words and non-words (Figure 1-2) influenced processing of the statistics-based phoneme features, which in turn may have indirectly affected the phoneme onset responses due to their concurrent timing. Additional activation of brain regions in the processing of non-words may also have contributed to the phoneme onset difference.

The temporal structure of non-words’ cohort entropy closely resembled the word onset responses, especially compared to those of phoneme surprisal. While one might expect similar trends between the phoneme surprisal and cohort entropy, it is important to note that these measures quantify different aspects of sub-lexical processing: phoneme surprisal represents *phonological* predictability, whereas cohort entropy reflects *lexical* uncertainty (Gwilliams and Davis, 2022). In this sense, cohort entropy effects likely reflect lexical processing more than just phoneme level processing, and this is supported by these results. Considerable temporal separation between early and middle peaks of word onset and cohort entropy may suggest additional mechanisms associated with non-word processing. A key difference between segmenting non-words vs. words is that boundaries between non-words are not clearly defined, and identifying them relies on indirect cues (e.g., prosody changes). When early and middle peaks were combined, no amplitude differences were observed between passage types, only latency, indicating that they indeed represent a single source that is linked to word segmentation (a bottom-up mechanism), but the latency of which depends upon the difficulty of the segmentation problem.

In general, the late peaks for phoneme and lexical level features were stronger in scrambled words compared to non-words and narrative passages, and were delayed from narrative to scrambled to non-words, suggesting the late responses are affected by linguistic content. Remarkably, this late peak resembles the characteristics of the N400 ERP response, a well-known brain response modulated by comprehension and predictability, often used to investigate semantic processing (Lau et al., 2008; Kutas and Federmeier, 2011). Additionally, the N400 plays a key role in predictive coding frameworks, where it has a natural and compelling interpretation as representing prediction error (Nour Eddine et al., 2024). Within the N400-like responses seen here at lexical and sub-lexical levels, our results demonstrate predictive coding operating at multiple stages. Our results suggest that context-based predictability facilitates the pre-activation of semantic integration, thereby reducing the strength and latency of N400-like response in narrative passages compared to scrambled words (Lam et al., 2016; Slaats et al., 2023). Some studies have also reported weaker late response with scrambling (Broderick et al., 2022; Gillis et al., 2023), though this difference may be related to the variations in the experimental design (EEG vs MEG, contextual measure employed). Conversely, the smaller N400-like responses for non-words aligns with non-words being mostly unpredictable, and thus not activating the N400 mechanism. However, in the current study the non-word passages did include some non-words that resembled real words (e.g., “sustument” and “bi”), which could lead to lexical activation of root words, and, consequently, elicit some N400 response. Some lexical activation for non-words could diminish the difference between narrative and non-words. Therefore, the N400-like response seen here could arise from both semantic and non-semantic violations of expectation. These interpretations are further supported by the latency analysis, which showed that peaks are delayed from narrative to scrambled to non-words. The earliest processing of the narrative stimulus suggests that rapid access to the mental lexicon is facilitated by the contextual information (Deacon et al., 2004; Kutas and Federmeier, 2011). Other studies have shown that the N400 is stronger for non-words compared to words (Bentin et al., 1999; Holcomb, 2007), however, but in paradigms where the non-words were presented between meaningful words, which alters the experimental design, behavioral expectations, and, likely, the neural processing form the current work. These results suggest that the late responses to both phoneme and lexical features are influenced by top-down driven mechanisms. While bottom-up driven mechanisms are less intriguing, top-down driven mechanisms demonstrate involvement in predictive coding mechanisms, making them better neural markers of cognitive processes.

## Acknowledgments

This work was supported by the National Institutes of Health grant R01-DC019394 and National Science Foundation grant SMA 1734892. We thank Vinoj Jayasundara for his assistance in context-based linguistic feature generation and Ciaran Stone for excellent technical assistance in MEG data recordings.

## Author Contributions

I.M.D.K. and J.Z.S. conceived and designed the experiments; I.M.D.K. collected data; I.M.D.K., C.B., S.B., P.R., and J.Z.S. designed the analysis, analyzed data and interpreted data; I.M.D.K. drafted manuscript; I.M.D.K., C.B., S.B., P.R., and J.Z.S. edited and revised manuscript.

## Extended Data

**Figure 1-1.**
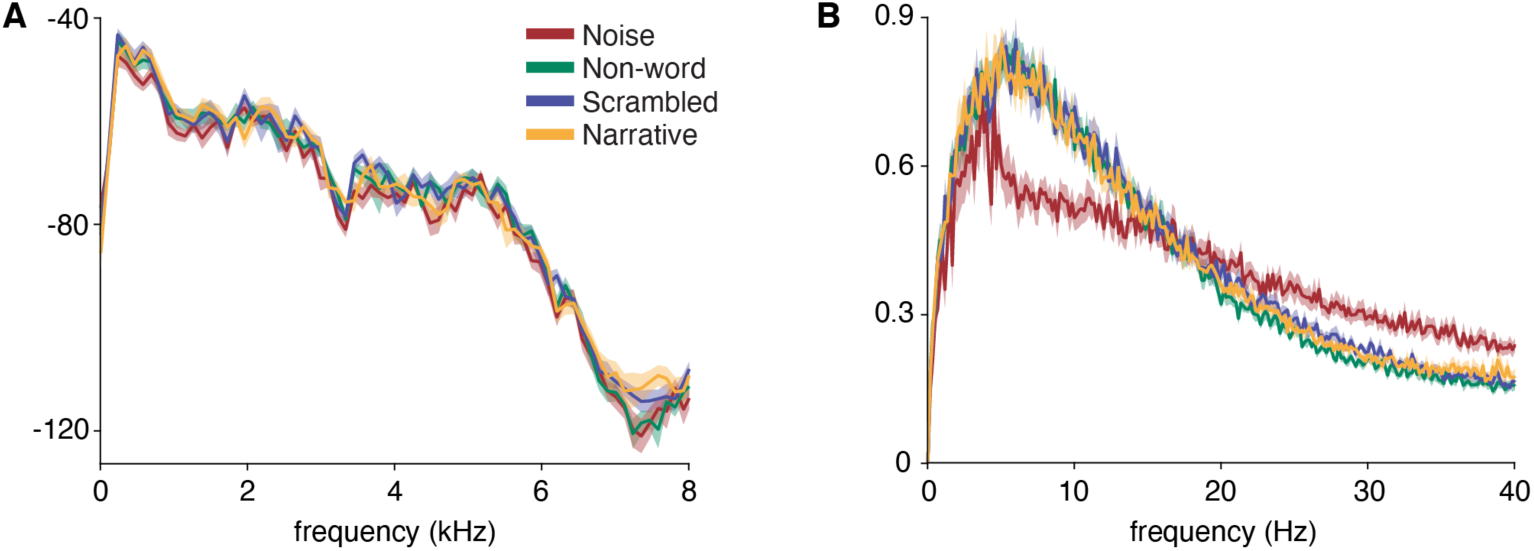
Comparison of Stimulus Acoustic Properties. (A). Periodograms and (B). Modulation spectrum obtained using the methods of Ding et al., (2017). Even though the spectral characteristics are similar between the stimulus types, the slow temporal modulation is different between speech and non-speech. There is no visible difference in acoustic properties between the speech passages. Periodograms and modulation spectra were computed for 10 chunks of 6 seconds each, per each passage type and then mean ± standard error is plotted to illustrate data.

**Figure 1-2.**
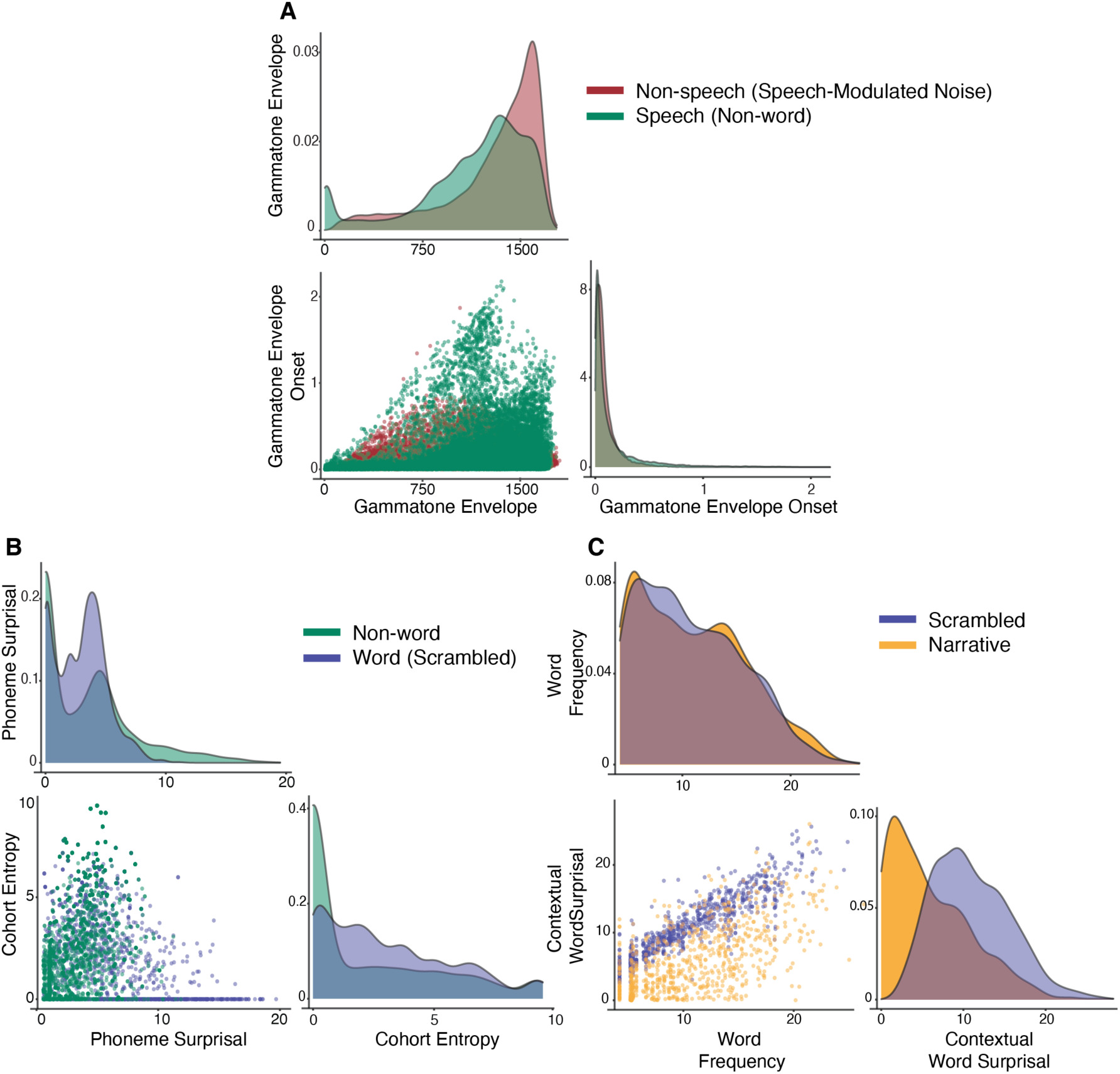
Comparison of predictor variables between passage types. (a). Acoustic feature comparisons between non-speech and (non-word) speech passage: they share similarities in the distribution of envelope onset predictor values, but not for envelope predictors. (b). Phoneme surprisal and cohort entropy comparison between non-words and meaningful words (scrambled passage): both predictor distributions depend strongly on the stimulus type. (c). Word frequency and contextual word surprisal comparisons between scrambled and narrative passages: the two word frequency distributions are nearly identical, by design, but the contextual word surprisal distributions diverge strongly (as expected, the narrative case is strongly biased toward low surprisal; additionally, in the scrambled word case, both forms of surprisal are highly correlated, collapsing into a narrow diagonal distribution. In each panel, the top and right plots show frequency histograms that present the distribution of each feature, where the y-axis represents the bin density of points, scaled to integrate to one.; the bottom left scatterplot shows a visualization of the correlation between the two predictor variables.

**Figure 2-1.**
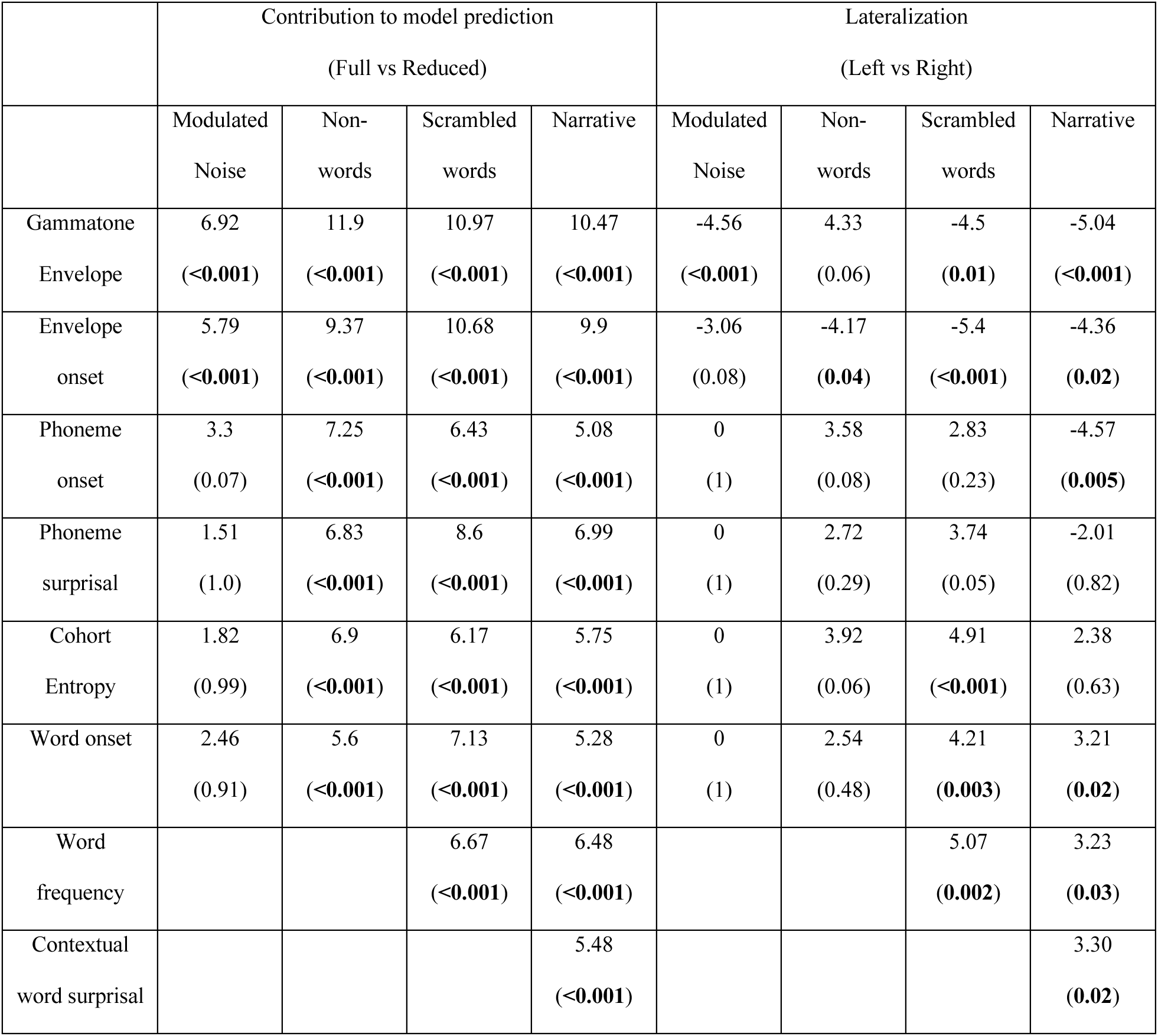
Summary statistics table for the model prediction comparisons. *t*_max_ and corresponding *p* values are reported. Second column summarizes the contribution of each feature to the model’s predictive power. Third column summarizes the lateralization results.

**Figure 3-1.**
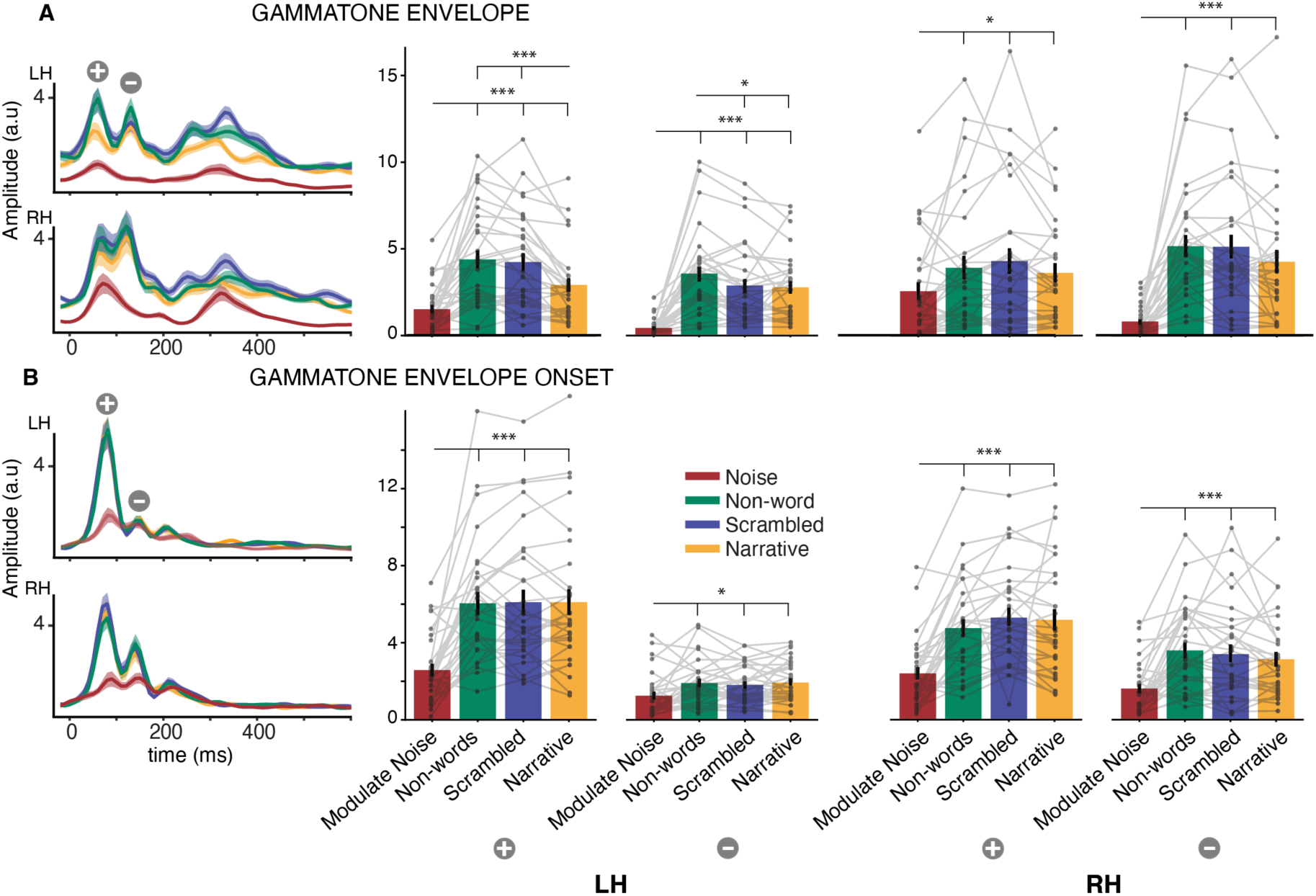
Neural Responses to acoustic features. (A) Gammatone envelope and (B) Gammatone envelope onset in left (LH) and right (RH) hemispheres. This figure expands on the data shown in Figure 3. The TRFs exhibit an early positive and a late negative polarity peak indicated by 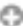 and 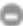 respectively. Right panel bar plots compare the peak amplitudes across passage types. LH and RH denotes left and right hemisphere respectively. Both early and late responses are stronger for speech compared to non-speech. Differences between the speech passages were found only for the envelope responses and in the left hemisphere. \**p*<0.05, \*\**p*<0.01, \*\*\**p*<0.001

**Figure 3-2.**
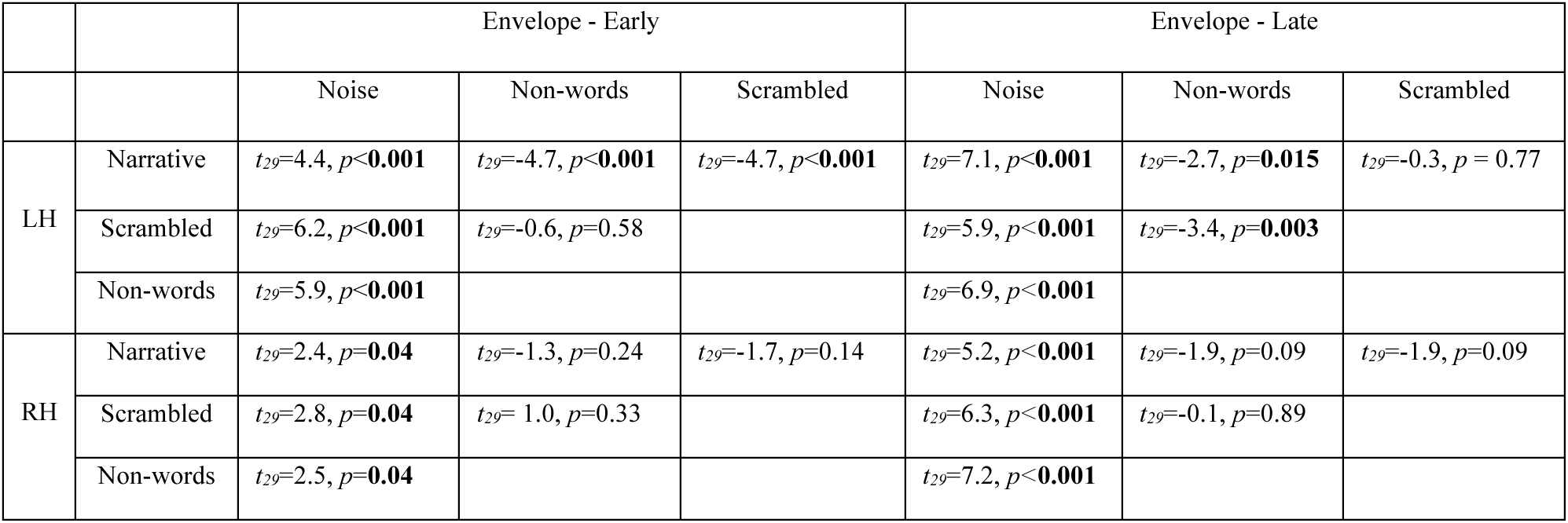
Summary Statistics table for envelope TRF peak amplitude comparisons. *P*-values are corrected for multiple comparisons using false discovery rate (FDR). LH and RH represent left and right hemispheres respectively.

**Figure 3-3.**
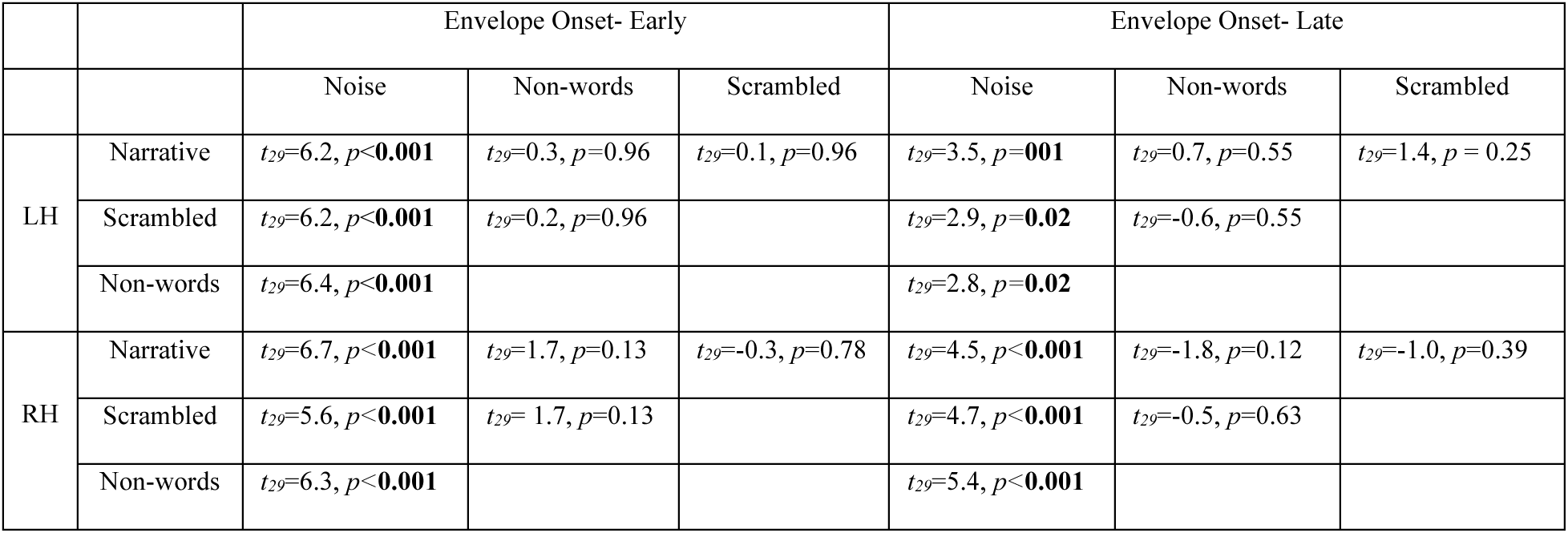
Summary Statistics table for envelope onset TRF peak amplitude comparisons. Other details as in Figure 3-2.

**Figure 4-1.**
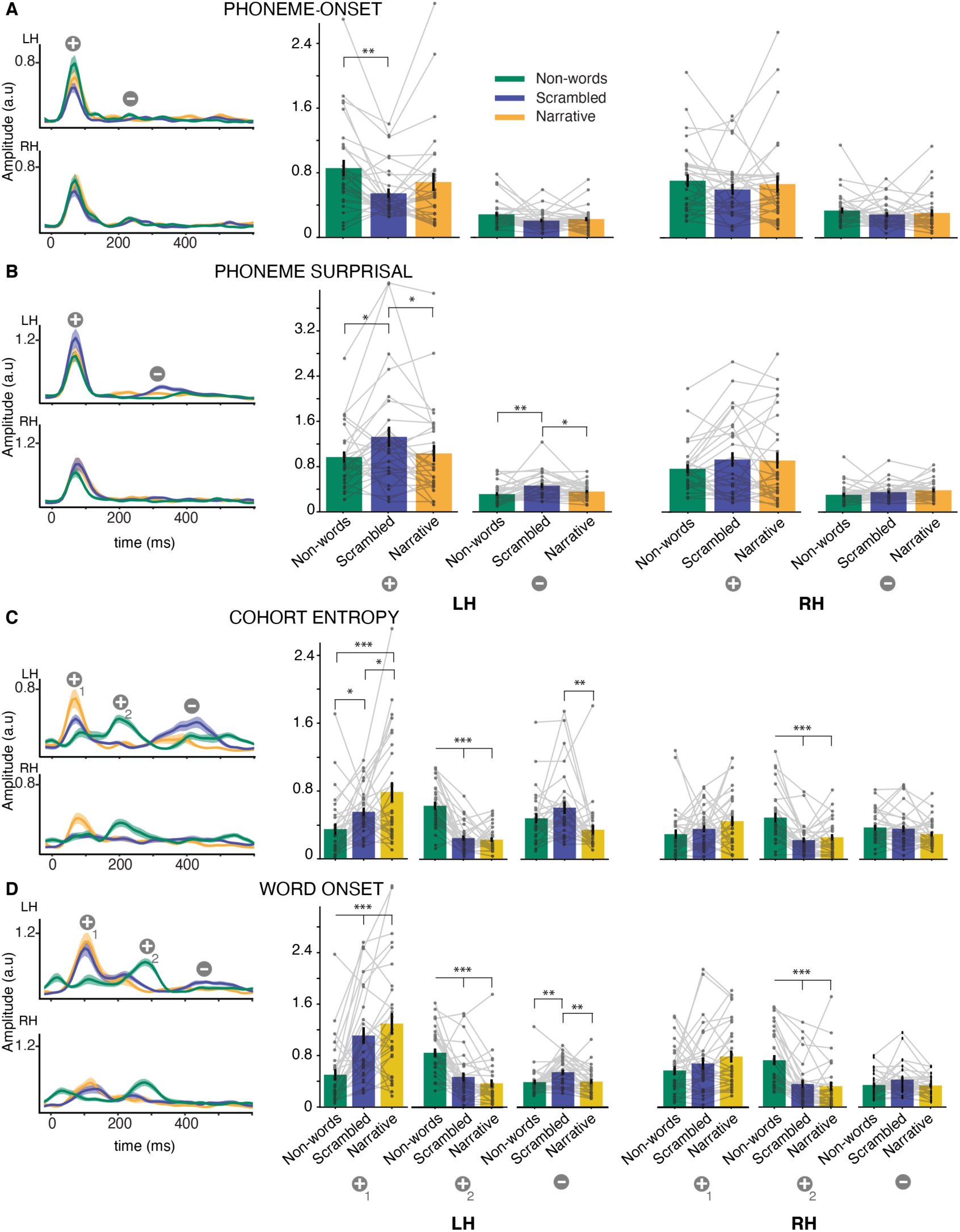
Neural responses to sub-lexical and word onset speech features. (A). Phoneme onset, (B). word onset, (C). phoneme surprisal, and (D). cohort entropy (TRF magnitude plots and TRF peak bar plots as in Figure 3-1). This figure expands on the data shown in Figure 4. TRFs exhibit an early positive and a late negative polarity peak indicated by 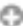 and 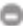 respectively. For both word onset and cohort entropy responses, non-words showed a robust positive polarity peak between early and late peaks. These early, middle, and late peaks are indicated by 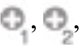 and 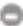respectively. The right column bar plots compare the peak amplitudes across passage types. LH and RH denotes left and right hemisphere respectively. Overall, the early responses were differently modulated by the linguistic content. The middle peak was stronger for non-words, while the late peak was stronger for scrambled passages. No differences, except the strong middle responses for non-words were found in the right hemisphere.

**Figure 4-2.**
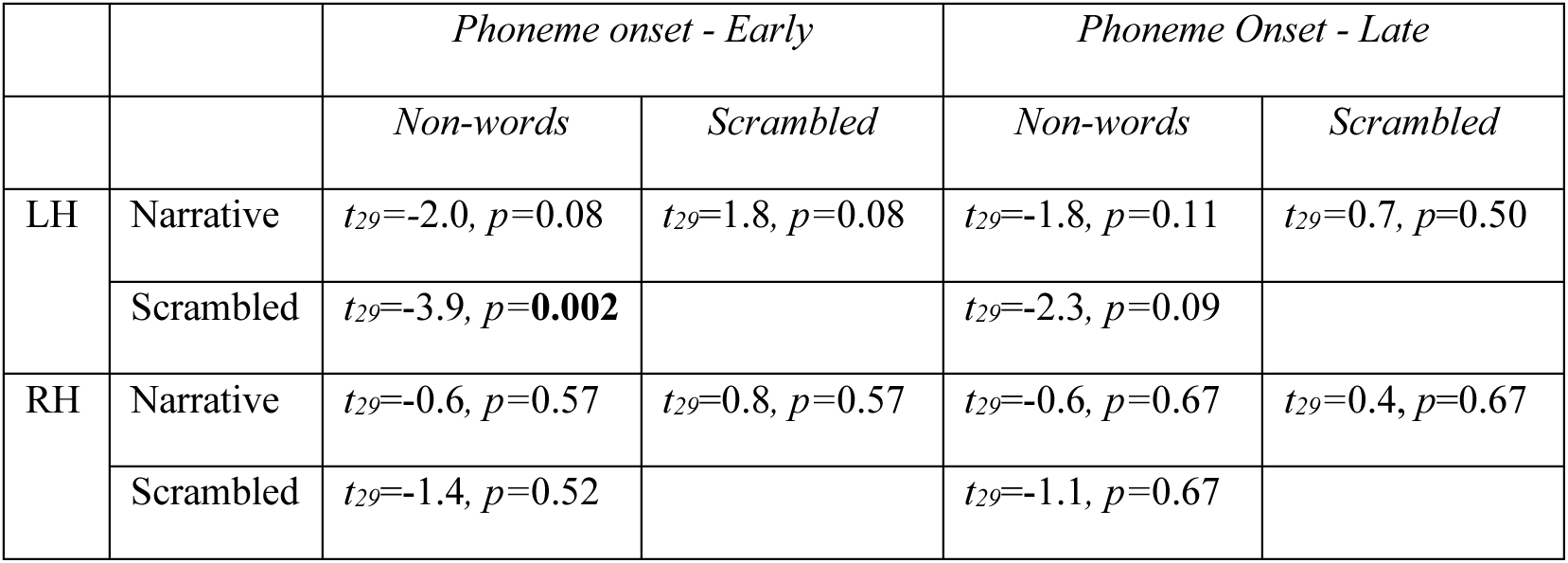
Summary Statistics table for phoneme onset TRF peak amplitude comparisons. Other details as in Figure 3-2.

**Figure 4-3.**
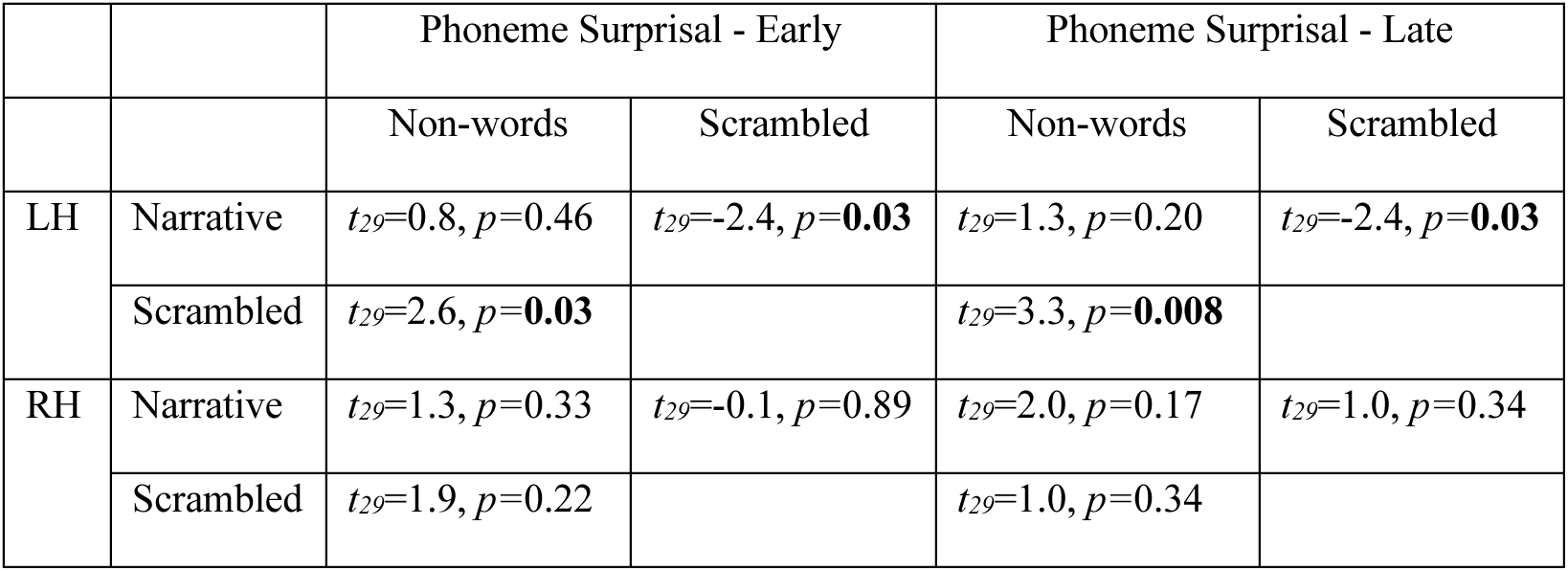
Summary Statistics table for phoneme surprisal TRF peak amplitude comparisons. Other details as in Figure 3-2.

**Figure 4-4.**
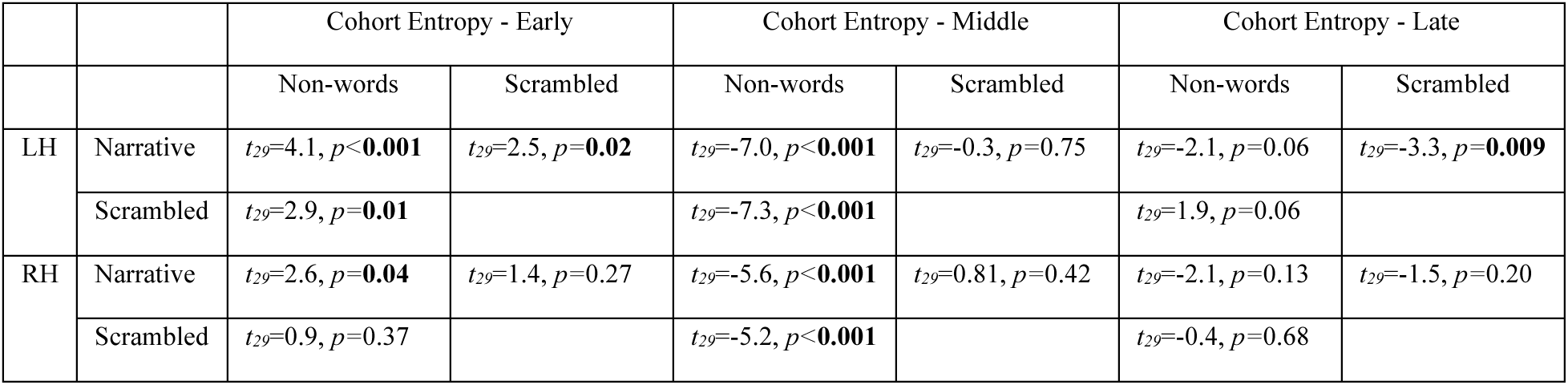
Summary Statistics table for cohort entropy TRF peak amplitude comparisons. Other details as in Figure 3-2.

**Figure 4-5.**
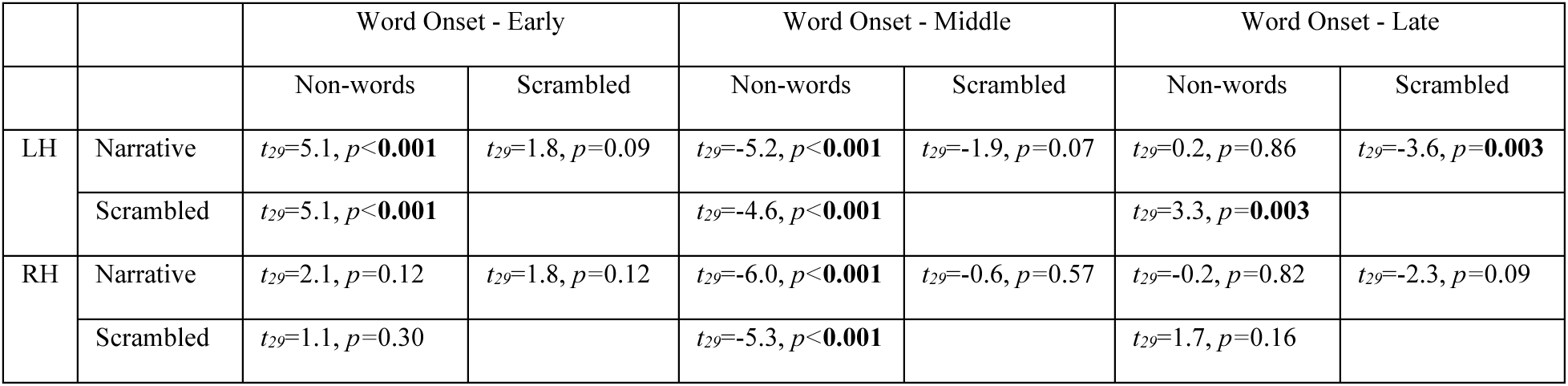
Summary Statistics table for word onset TRF peak amplitude comparisons. Other details as in Figure 3-2.

**Figure 4-6.**
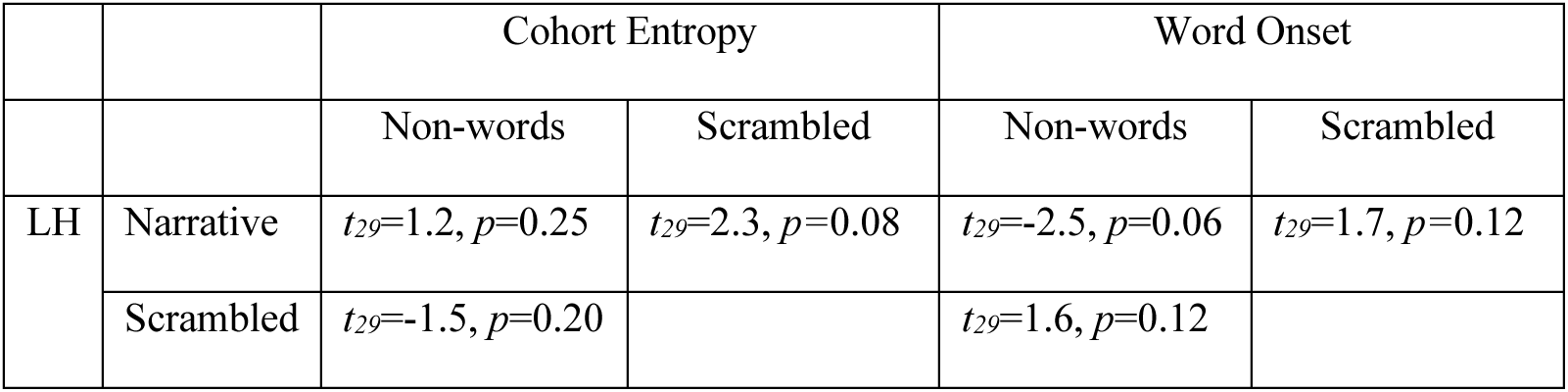
Summary statistics table for combined early and middle peak amplitude comparisons for cohort entropy and word onsets. Other details as in Figure 3-2.

**Figure 5-1.**
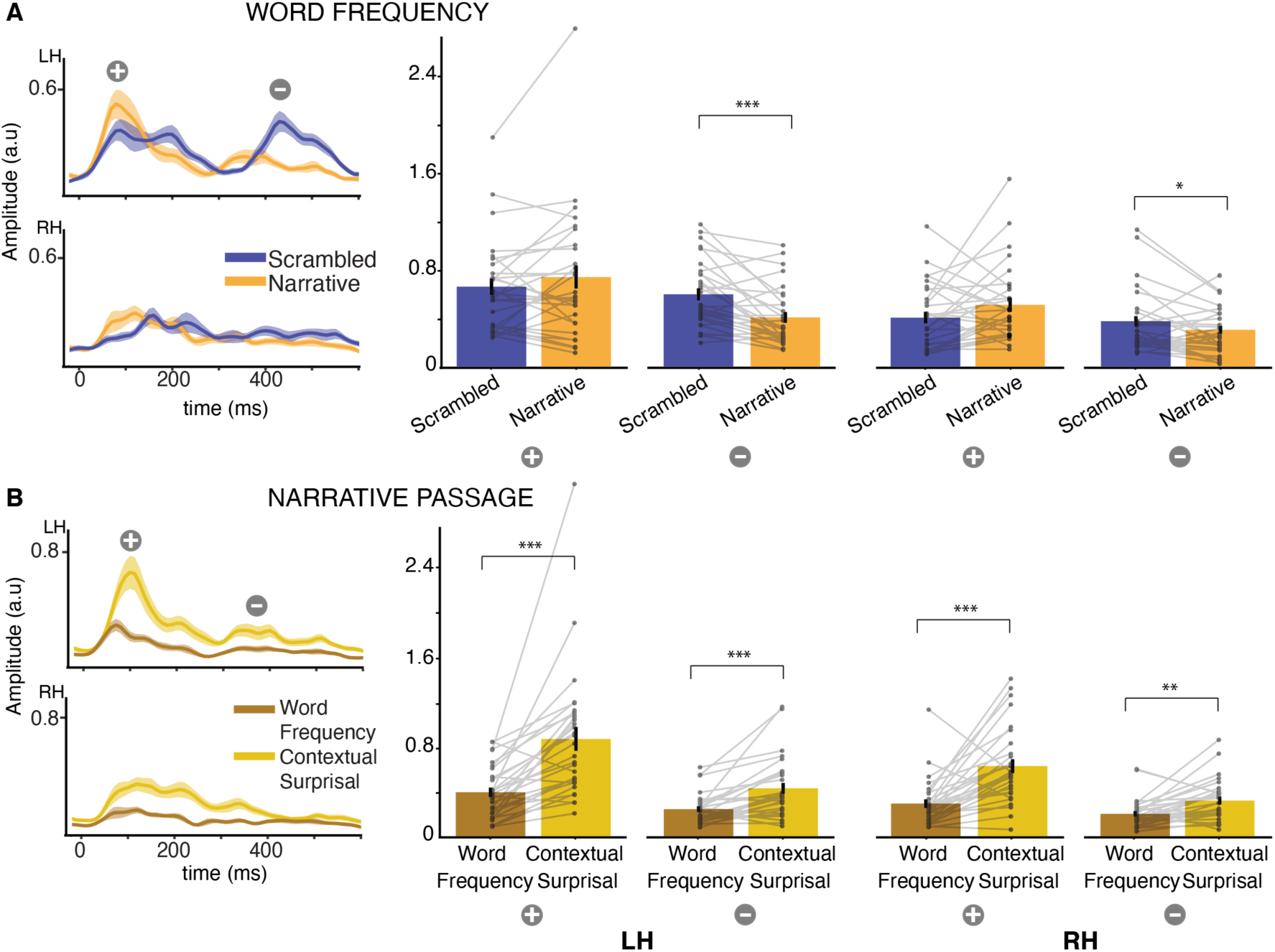
Neural responses to lexico-semantic features. (A) Word frequency and (B) Word frequency vs contextual word surprisal for the narrative passage (TRF magnitude plots and TRF peak bar plots as in Figure 3-1). This figure expands on the data shown in Figure 5. The late peak in word frequency responses is stronger for scrambled words compared to narrative passages. Contextual word surprisal is stronger compared to local word surprisal (word frequency). LH and RH denotes left and right hemisphere respectively. TRFs exhibit an early positive and a late negative polarity peak indicated by 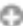 and 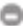 respectively. The late word frequency responses (N400-like) are stronger for scrambled passages compared to narrative passage. Contextual word surprisal responses are stronger compared to word frequency responses. Note that the peak amplitudes for word frequency in (A) and (B) are different, as the TRF model in (A) does not include contextual surprisal.

**Figure 5-2.**
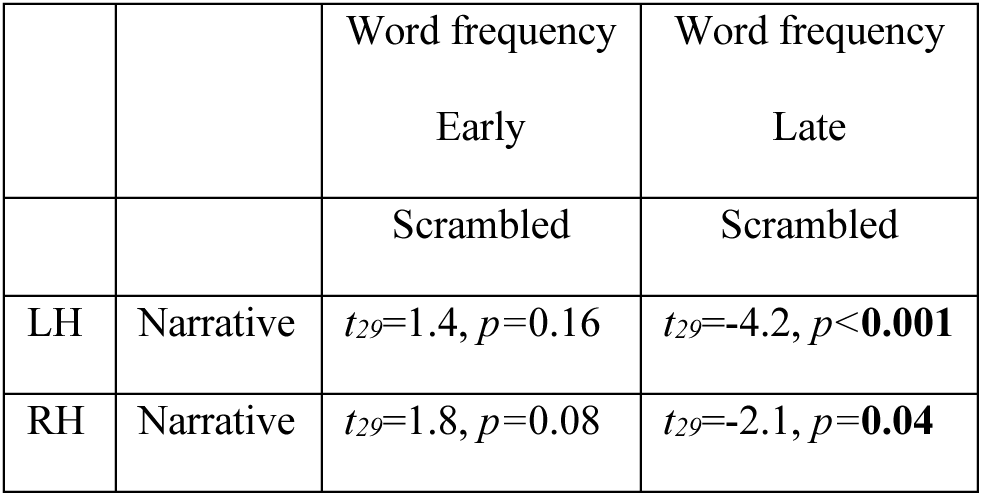
Summary Statistics table for word frequency TRF peak amplitude comparisons. Other details as in Figure 3-2.

**Figure 5-3.**
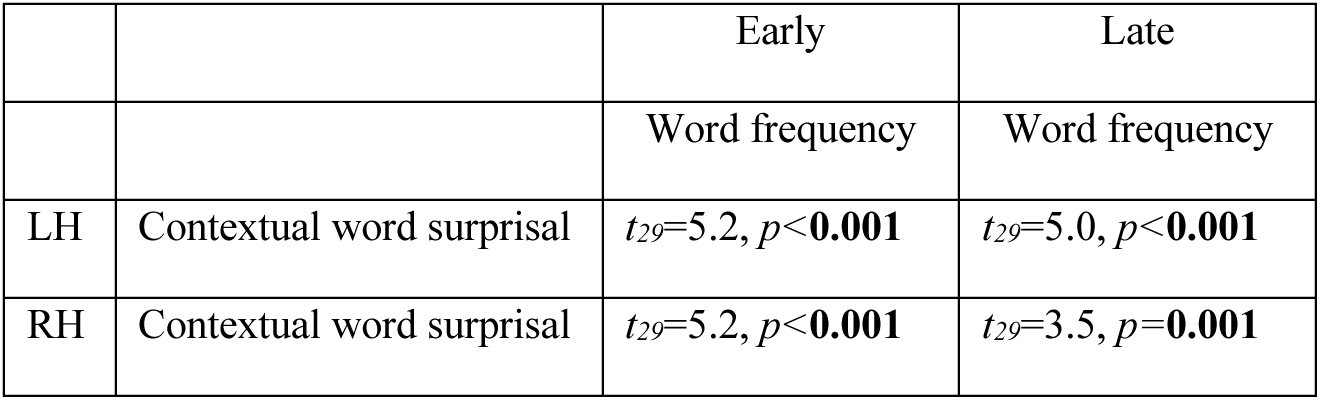
Summary Statistics table for contextual surprisal vs word frequency TRF peak amplitude comparisons. Other details as in Figure 3-2.

## References

Aiken SJ, Picton TW (2008) Human Cortical Responses to the Speech Envelope. Ear & Hearing 29:139– 157.

Alday PM (2019) M/EEG analysis of naturalistic stories: a review from speech to language processing. Language, Cognition and Neuroscience 34:457–473

Aoki NB, Cohn M, Zellou G (2022) The clear speech intelligibility benefit for text-to-speech voices: Effects of speaking style and visual guise. JASA Express Letters 2:045204.

Arnal LH, Poeppel D, Giraud A-L (2016) A Neurophysiological Perspective on Speech Processing in “The Neurobiology of Language.” In: Neurobiology of Language, pp 463–478. Elsevier.

Bell AJ, Sejnowski TJ (1995) An Information-Maximization Approach to Blind Separation and Blind Deconvolution. Neural Computation 7:1129–1159.

Bentin S, Mouchetant-Rostaing Y, Giard MH, Echallier JF, Pernier J (1999) ERP Manifestations of Processing Printed Words at Different Psycholinguistic Levels: Time Course and Scalp Distribution. Journal of Cognitive Neuroscience 11:235–260.

Binder JR (2000) Human Temporal Lobe Activation by Speech and Nonspeech Sounds. Cerebral Cortex 10:512–528.

Boersma P, Weenink D (2021) Praat: doing phonetics by computer.

Bozic M, Tyler LK, Ives DT, Randall B, Marslen-Wilson WD (2010) Bihemispheric foundations for human speech comprehension. Proc Natl Acad Sci USA 107:17439–17444.

Bradshaw AR, Thompson PA, Wilson AC, Bishop DVM, Woodhead ZVJ (2017) Measuring language lateralisation with different language tasks: a systematic review. PeerJ 5:e3929.

Brodbeck C, Bhattasali S, Cruz Heredia AA, Resnik P, Simon JZ, Lau E (2022) Parallel processing in speech perception with local and global representations of linguistic context. eLife 11:e72056.

Brodbeck C, Das P, Gillis M, Kulasingham JP, Bhattasali S, Gaston P, Resnik P, Simon JZ (2023) Eelbrain, a Python toolkit for time-continuous analysis with temporal response functions. eLife 12:e85012.

Brodbeck C, Hong LE, Simon JZ (2018) Rapid Transformation from Auditory to Linguistic Representations of Continuous Speech. Current Biology 28:3976–3983.e5.

Brodbeck C, Jiao A, Hong LE, Simon JZ (2020) Neural speech restoration at the cocktail party: Auditory cortex recovers masked speech of both attended and ignored speakers Malmierca MS, ed. PLoS Biol 18:e3000883.

Broderick MP, Zuk NJ, Anderson AJ, Lalor EC (2022) More than words: Neurophysiological correlates of semantic dissimilarity depend on comprehension of the speech narrative. Eur J of Neuroscience 56:5201–5214.

Caucheteux C, Gramfort A, King J-R (2023) Evidence of a predictive coding hierarchy in the human brain listening to speech. Nat Hum Behav 7:430–441.

Dale AM, Sereno MI (1993) Improved Localizadon of Cortical Activity by Combining EEG and MEG with MRI Cortical Surface Reconstruction: A Linear Approach. Journal of Cognitive Neuroscience 5:162–176.

Dambacher M, Kliegl R, Hofmann M, Jacobs AM (2006) Frequency and predictability effects on event-related potentials during reading. Brain Research 1084:89–103.

David SV, Mesgarani N, Shamma SA (2007) Estimating sparse spectro-temporal receptive fields with natural stimuli. Network: Computation in Neural Systems 18:191–212.

Davis MH, Ford MA, Kherif F, Johnsrude IS (2011) Does Semantic Context Benefit Speech Understanding through “Top–Down” Processes? Evidence from Time-resolved Sparse fMRI. Journal of Cognitive Neuroscience 23:3914–3932.

De Cheveigné A (2024) Predictive coding in the auditory brainstem. Neuroscience.

Deacon D, Dynowska A, Ritter W, Grose-Fifer J (2004) Repetition and semantic priming of nonwords: Implications for theories of N400 and word recognition. Psychophysiology 41:60–74.

Deniz F, Tseng C, Wehbe L, Dupré La Tour T, Gallant JL (2023) Semantic Representations during Language Comprehension Are Affected by Context. J Neurosci 43:3144–3158.

Desikan RS, Ségonne F, Fischl B, Quinn BT, Dickerson BC, Blacker D, Buckner RL, Dale AM, Maguire RP, Hyman BT, Albert MS, Killiany RJ (2006) An automated labeling system for subdividing the human cerebral cortex on MRI scans into gyral based regions of interest. Neuroimage 31:968–980.

Di Liberto GM, O’Sullivan JA, Lalor EC (2015) Low-Frequency Cortical Entrainment to Speech Reflects Phoneme-Level Processing. Current Biology 25:2457–2465.

Ding N, Patel AD, Chen L, Butler H, Luo C, Poeppel D (2017) Temporal modulations in speech and music. Neuroscience & Biobehavioral Reviews 81:181–187.

Ding N, Simon JZ (2012a) Neural coding of continuous speech in auditory cortex during monaural and dichotic listening. Journal of Neurophysiology 107:78–89.

Ding N, Simon JZ (2012b) Emergence of neural encoding of auditory objects while listening to competing speakers. Proceedings of the National Academy of Sciences 109:11854–11859.

Fedorenko E, Nieto-Castañon A, Kanwisher N (2012) Lexical and syntactic representations in the brain: An fMRI investigation with multi-voxel pattern analyses. Neuropsychologia 50:499–513.

Fiedler L, Wöstmann M, Herbst SK, Obleser J (2019) Late cortical tracking of ignored speech facilitates neural selectivity in acoustically challenging conditions. NeuroImage 186:33–42.

Fischl B (2012) FreeSurfer. NeuroImage 62:774–781.

Fishbach A, Nelken I, Yeshurun Y (2001) Auditory Edge Detection: A Neural Model for Physiological and Psychoacoustical Responses to Amplitude Transients. Journal of Neurophysiology 85:2303–2323.

Gaston P, Brodbeck C, Phillips C, Lau E (2023) Auditory Word Comprehension Is Less Incremental in Isolated Words. Neurobiology of Language 4:29–52.

Gillis M, Vanthornhout J, Francart T (2023) Heard or Understood? Neural Tracking of Language Features in a Comprehensible Story, an Incomprehensible Story and a Word List. eNeuro 10:ENEURO.0075-23.2023.

Gillis M, Vanthornhout J, Simon JZ, Francart T, Brodbeck C (2021) Neural Markers of Speech Comprehension: Measuring EEG Tracking of Linguistic Speech Representations, Controlling the Speech Acoustics. J Neurosci 41:10316–10329.

Gow DW (2012) The cortical organization of lexical knowledge: A dual lexicon model of spoken language processing. Brain and Language 121:273–288.

Gramfort A (2013) MEG and EEG data analysis with MNE-Python. Front Neurosci 7.

Gramfort A, Luessi M, Larson E, Engemann DA, Strohmeier D, Brodbeck C, Parkkonen L, Hämäläinen MS (2014) MNE software for processing MEG and EEG data. NeuroImage 86:446–460.

Gwilliams L, Davis MH (2022) Extracting Language Content from Speech Sounds: The Information Theoretic Approach. In: Speech Perception (Holt LL, Peelle JE, Coffin AB, Popper AN, Fay RR, eds), pp 113–139 Springer Handbook of Auditory Research. Cham: Springer International Publishing.

Gwilliams L, King J-R, Marantz A, Poeppel D (2022) Neural dynamics of phoneme sequences reveal position-invariant code for content and order. Nat Commun 13:6606.

Gwilliams L, Marantz A (2015) Non-linear processing of a linear speech stream: The influence of morphological structure on the recognition of spoken Arabic words. Brain and Language 147:1– 13.

Hämäläinen MS, Ilmoniemi RJ (1994) Interpreting magnetic fields of the brain: minimum norm estimates. Med Biol Eng Comput 32:35–42.

Hamilton LS, Edwards E, Chang EF (2018) A Spatial Map of Onset and Sustained Responses to Speech in the Human Superior Temporal Gyrus. Current Biology 28:1860–1871.e4.

Heeris J (2018) Gammatone Filterbank Toolkit.

Heilbron M, Armeni K, Schoffelen J-M, Hagoort P, de Lange FP (2022) A hierarchy of linguistic predictions during natural language comprehension. Proc Natl Acad Sci USA 119:e2201968119.

Herrmann B (2023) The perception of artificial-intelligence (AI) based synthesized speech in younger and older adults. Int J Speech Technol.

Hickok G, Poeppel D (2004) Dorsal and ventral streams: a framework for understanding aspects of the functional anatomy of language. Cognition 92:67–99.

Hickok G, Poeppel D (2007) The cortical organization of speech processing. Nat Rev Neurosci 8:393–402.

Holcomb PJ (2007) Semantic priming and stimulus degradation: Implications for the role of the N400 in language processing. Psychophysiology 30:47–61.

Huizeling E, Arana S, Hagoort P, Schoffelen J-M (2022) Lexical Frequency and Sentence Context Influence the Brain’s Response to Single Words. Neurobiology of Language 3:149–179.

Karunathilake IMD, Kulasingham JP, Simon JZ (2023) Neural tracking measures of speech intelligibility: Manipulating intelligibility while keeping acoustics unchanged. Proc Natl Acad Sci USA 120:e2309166120.

Keshishian M, Akkol S, Herrero J, Bickel S, Mehta AD, Mesgarani N (2023) Joint, distributed and hierarchically organized encoding of linguistic features in the human auditory cortex. Nat Hum Behav 7:740–753.

Kubanek J, Brunner P, Gunduz A, Poeppel D, Schalk G (2013) The Tracking of Speech Envelope in the Human Cortex Rodriguez-Fornells A, ed. PLoS ONE 8:e53398.

Kutas M, Federmeier KD (2011) Thirty Years and Counting: Finding Meaning in the N400 Component of the Event-Related Brain Potential (ERP). Annu Rev Psychol 62:621–647.

Lalor EC, Power AJ, Reilly RB, Foxe JJ (2009) Resolving Precise Temporal Processing Properties of the Auditory System Using Continuous Stimuli. Journal of Neurophysiology 102:349–359.

Lam NHL, Schoffelen J-M, Uddén J, Hultén A, Hagoort P (2016) Neural activity during sentence processing as reflected in theta, alpha, beta, and gamma oscillations. NeuroImage 142:43–54.

Lau EF, Phillips C, Poeppel D (2008) A cortical network for semantics: (de)constructing the N400. Nat Rev Neurosci 9:920–933.

Lerner Y, Honey CJ, Silbert LJ, Hasson U (2011) Topographic Mapping of a Hierarchy of Temporal Receptive Windows Using a Narrated Story. J Neurosci 31:2906–2915.

Mai G, Minett JW, Wang WS-Y (2016) Delta, theta, beta, and gamma brain oscillations index levels of auditory sentence processing. NeuroImage 133:516–528.

McAuliffe M, Socolof M, Mihuc S, Wagner M, Sonderegger M (2017) Montreal Forced Aligner: Trainable Text-Speech Alignment Using Kaldi. In: Interspeech 2017, pp 498–502. ISCA.

Nour Eddine S, Brothers T, Wang L, Spratling M, Kuperberg GR (2024) A predictive coding model of the N400. Cognition 246:105755.

Nourski KV, Steinschneider M, Rhone AE, Kovach CK, Kawasaki H, Howard MA (2019) Differential responses to spectrally degraded speech within human auditory cortex: An intracranial electrophysiology study. Hearing Research 371:53–65.

Oganian Y, Chang EF (2019) A speech envelope landmark for syllable encoding in human superior temporal gyrus. Sci Adv 5:eaay6279.

Oord A van den, Dieleman S, Zen H, Simonyan K, Vinyals O, Graves A, Kalchbrenner N, Senior A, Kavukcuoglu K (2016) WaveNet: A Generative Model for Raw Audio. In: Proc. 9th ISCA Workshop on Speech Synthesis Workshop (SSW 9), pp 125.

O’Sullivan JA, Power AJ, Mesgarani N, Rajaram S, Foxe JJ, Shinn-Cunningham BG, Slaney M, Shamma SA, Lalor EC (2015) Attentional Selection in a Cocktail Party Environment Can Be Decoded from Single-Trial EEG. Cerebral Cortex 25:1697–1706.

Overath T, McDermott JH, Zarate JM, Poeppel D (2015) The cortical analysis of speech-specific temporal structure revealed by responses to sound quilts. Nat Neurosci 18:903–911.

Overath T, Paik JH (2021) From acoustic to linguistic analysis of temporal speech structure: Acousto-linguistic transformation during speech perception using speech quilts. NeuroImage 235:117887.

Payne BR, Lee C-L, Federmeier KD (2015) Revisiting the incremental effects of context on word processing: Evidence from single-word event-related brain potentials: Effects of context on world-level N400. Psychophysiol 52:1456–1469.

Peelle JE (2012) The hemispheric lateralization of speech processing depends on what “speech” is: a hierarchical perspective. Front Hum Neurosci 6.

Poeppel D (2003) The analysis of speech in different temporal integration windows: cerebral lateralization as ‘asymmetric sampling in time.’ Speech Communication 41:245–255.

Pollan M (2001) The botany of desire : a plant’s eye view of the world, 1st ed. New York: Random House.

Price CJ (2012) A review and synthesis of the first 20years of PET and fMRI studies of heard speech, spoken language and reading. NeuroImage 62:816–847.

R Core Team (2020) R: A Language and Environment for Statistical Computing.

Ross ED, Thompson RD, Yenkosky J (1997) Lateralization of Affective Prosody in Brain and the Callosal Integration of Hemispheric Language Functions. Brain and Language 56:27–54.

Schrimpf M, Blank IA, Tuckute G, Kauf C, Hosseini EA, Kanwisher N, Tenenbaum JB, Fedorenko E (2021) The neural architecture of language: Integrative modeling converges on predictive processing. Proc Natl Acad Sci USA 118:e2105646118.

Sennrich R, Haddow B, Birch A (2016) Neural Machine Translation of Rare Words with Subword Units. In: Proceedings of the 54th Annual Meeting of the Association for Computational Linguistics (Volume 1: Long Papers), pp 1715–1725. Berlin, Germany: Association for Computational Linguistics.

Shuai L, Gong T (2014) Temporal relation between top-down and bottom-up processing in lexical tone perception. Front Behav Neurosci.

Slaats S, Weissbart H, Schoffelen J-M, Meyer AS, Martin AE (2023) Delta-Band Neural Responses to Individual Words Are Modulated by Sentence Processing. J Neurosci 43:4867–4883.

Smith S, Nichols T (2009) Threshold-free cluster enhancement: Addressing problems of smoothing, threshold dependence and localisation in cluster inference. NeuroImage 44:83–98.

Speer R, Chin J, Lin A, Jewett S, Nathan L (2018) LuminosoInsight/wordfreq: v2.2.

Steinschneider M, Nourski KV, Fishman YI (2013) Representation of speech in human auditory cortex: Is it special? Hearing Research 305:57–73.

Taulu S, Simola J (2006) Spatiotemporal signal space separation method for rejecting nearby interference in MEG measurements. Phys Med Biol 51:1759–1768.

Tezcan F, Weissbart H, Martin AE (2023) A tradeoff between acoustic and linguistic feature encoding in spoken language comprehension. eLife 12:e82386.

Vaswani A, Shazeer N, Parmar N, Uszkoreit J, Jones L, Gomez AN, Kaiser Ł, Polosukhin I (2017) Attention is All you Need. In: Advances in Neural Information Processing Systems (Guyon I, Luxburg UV, Bengio S, Wallach H, Fergus R, Vishwanathan S, Garnett R, eds). Curran Associates, Inc.

Wolf T et al. (2020) Transformers: State-of-the-Art Natural Language Processing. In: Proceedings of the 2020 Conference on Empirical Methods in Natural Language Processing: System Demonstrations, pp 38–45. Online: Association for Computational Linguistics.

Xu J, Kemeny S, Park G, Frattali C, Braun A (2005) Language in context: emergent features of word, sentence, and narrative comprehension. NeuroImage 25:1002–1015.

